# The Small β-barrel Domain: A Survey-based Structural Analysis

**DOI:** 10.1101/140376

**Authors:** Philippe Youkharibache, Stella Veretnik, Qingliang Li, Kimberly A. Stanek, Cameron Mura, Philip E. Bourne

## Abstract

The small β-barrel is an ancient protein structural domain characterized by extremes: It features an extremely broad range of structural varieties, a deeply intricate evolutionary history, and it is associated with a bewildering array of biomolecular pathways and physiological functions. These and related features of this domain are described and analyzed herein. Specifically, we present a comprehensive, survey-based analysis of the structural properties of small β-barrels (SBBs). We first consider the defining characteristics of the SBB fold, as well as the various systems of nomenclature used to describe it. In order to begin elucidating how such vast functional diversity is achieved by a relatively simple protein domain, we then explore the anatomy of the SBB fold and some of its representative structural variants. Many types of SBB proteins assemble into cyclic oligomers that act as the biologically-functional entity. These oligomers exhibit a great deal of plasticity even at the quaternary structural level—including homomeric and heteromeric assemblies, rings of variable subunit stoichiometries (pentamer, hexamer, etc.), as well as higher-order oligomers (e.g., double-rings) and fibrillar polymers. We conclude with three themes that emerge from the SBB’s unique structure↔function versatility.

## 1. Introduction

### 1.1 What is the small β-barrel (SBB) domain? Why study it?

The small β-barrel (SBB) is a phylogenetically pervasive and functionally diverse protein structural domain that we define and systematically analyze herein. SBBs, which comprise a subset of all β-barrel–containing protein domains, exhibit a complicated, richly intricate evolutionary history, and they are associated with myriad physiological functions. Apart from its β-rich secondary structural composition, the SBB domain is characterized by a small size (☐100 residues in five β-strands) and a highly conserved structural framework; as a reference point, membrane protein β-barrels range from eight to >20 strands, and even the smallest known ones exceed ≈150 residues, while larger ones are >700 residues (see Tables 1 in (Tamm, Hong, and Liang 2004; Fairman, Noinaj, and Buchanan 2011)). Another hallmark of the large set of known SBB-containing proteins is its unusually broad functional diversity. This diversity ranges, for instance, from (i) the often non-specific single-and double-stranded DNA-binding by an SBB known as the OB fold (Mitton-Fry et al. 2004), to (ii) RNA splicing, processing and other nucleic acid metabolic pathways (Vogel and Luisi 2011; Mura et al. 2013), to (iii) small noncoding RNA-based regulatory circuits (Vogel and Luisi 2011), to (iv) the translational machinery, including many ribosomal SBB proteins (Klein, Moore, and Steitz 2004; Valle et al. 2002), to (v) signal transduction and epigenetic regulation pathways (McCarty 1998; Patel and Wang 2013), to (vi) structural scaffolding of chromatin DNA in archaea (Robinson et al. 1998). The SBB structures found in these functional contexts are described in this text, and further information about biochemical functionalities can be found in the cited reviews. Note that the term ‘small β-barrel’ has appeared infrequently and sporadically in the literature over the past two decades. To our knowledge, no precise definition of an ‘SBB’ has been given; also, in such cases where the term has been used, the particular 3D structures either comply with our definition (given below), correspond to a subset of our definition, or else refer to *relatively* small membrane barrels (which still significantly exceed what is defined here as a ‘small β-barrel’).

Though their structures superficially resemble the β-barrels found in membrane proteins (many of which are functionally constrained, e.g., as transporters of small molecules through a central lumen), SBBs are significantly smaller in size, consisting of five or six short β-strands; these strands are often arranged as two closely-packed, nearly-orthogonal β-sheets (Figure 1, and detailed below). SBBs are highly flexible, in terms of their ligand-and substrate-binding capacities and, consequently, SBB-containing proteins are found in a broad range of cellular pathways—often as modules that bind, either specifically or generically, to various RNAs, DNAs and proteins. Despite its small size and relatively limited surface area, it appears that virtually every solvent-exposed region of an SBB can be adapted for binding to other biomolecules, depending on the functional context. In this regard, SBBs are reminiscent of the RNA recognition motif (RRM; Pfam clan CL0221)—a recurrent protein domain that binds both nucleic acids and proteins via many distinct ligand-binding structural motifs and biochemically-active surface patches (Cléry, Blatter, and Allain 2008).

**Figure 1.**
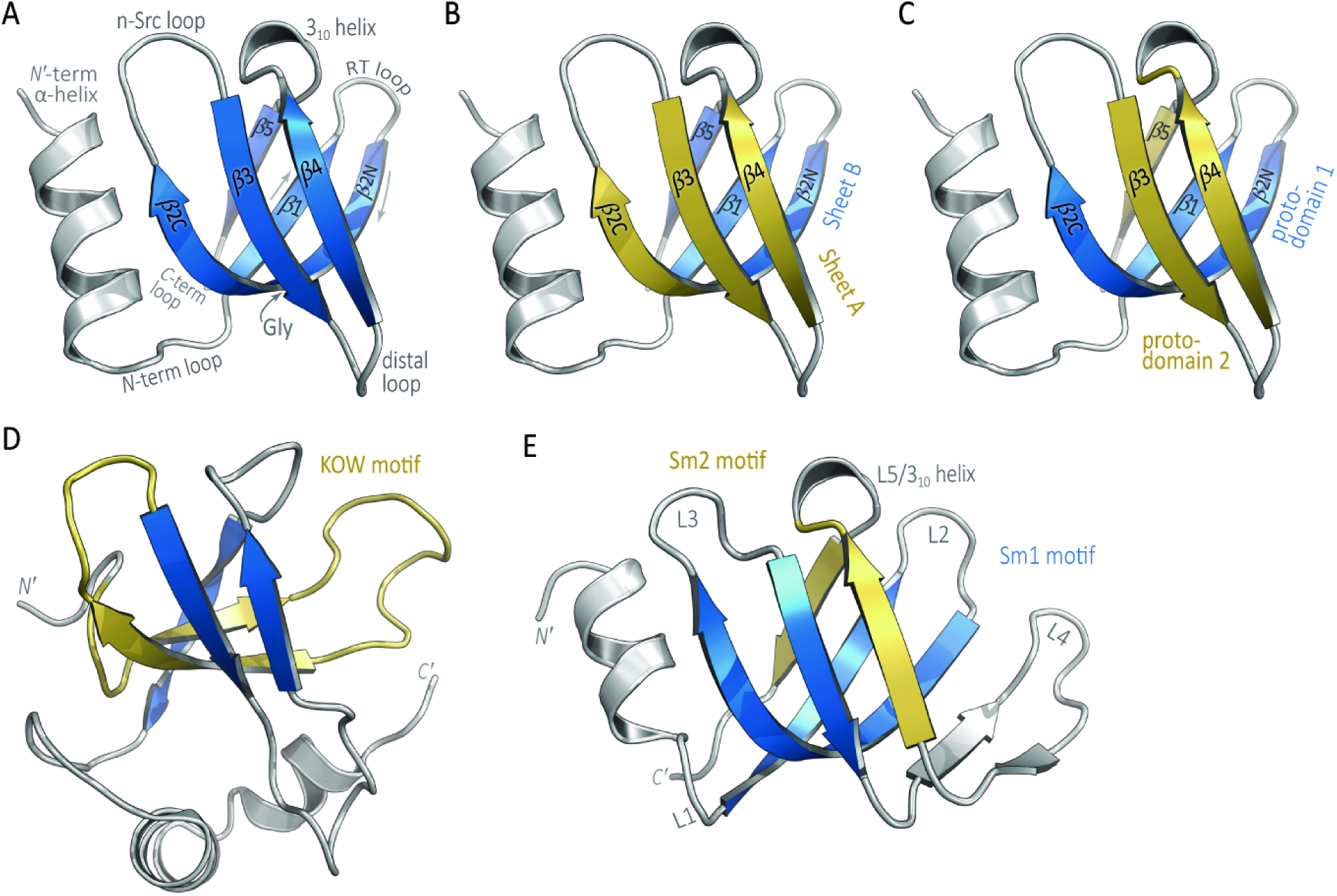
Structural overview and terminological features of the SBB domain. Hfq, a bacterial Sm-like protein (SCOP fold b.38.1.2), is taken as a reference structure throughout this work. Panel (A) shows the structure of S. *aureus* Hfq, drawn from the hexamer structure (PDB 1KQ2), with the SH3 strand numbering and loop terminology that we use for all SH3-like SBBs. The termini and other structural landmarks are labelled, including a conserved Gly near the point of greatest curvature in β2. In (B), the barrel is colored as two distinct β-sheets: *Sheet A* (*Meander*, yellow), which consists of β2C, β3 and β4, and *Sheet B (N-C;* blue), consisting of β5, β1 and β2N. In (C), the barrel is divided into two ‘proto-domains’ that are related via approximate C2 symmetry. Proto-domain 1 (blue) consists of strands β1, β2N and β2C; proto-domain 2 (yellow) consists of strands β3, β4 and β5. The KOW motif, shown in (D), consists of 27 residues from β1, β2, and the N’-term loop preceding β1. In the Sm fold, shown in (E) for the SmD3 protein, the β-sheet is often viewed as consisting of two functional motifs: strands β1→β3 comprise an ‘Sm1’ signature (wherein lie many residues involved in RNA-binding), and segment β4→β5 comprises an ‘Sm2’ motif (facilitates oligomerization, via these two edge strands). The loops in (D) are labeled using the Sm nomenclature (Table 1 in Section 2.3.1).

Unlike RRMs, many SBB-containing proteins exhibit a strong tendency to form toroidal discs and other higher-order structures that interact with various proteins or nucleic acids to serve as the biological functional unit. A notable example whereby SBB-mediated oligomerization yields an expanded range of biological functionality is the Sm/LSm class of RNA-associated proteins; the SBB domains of these proteins, found in Archaea, Bacteria and Eukarya, form a rich variety of homo-and hetero-oligomers. As detailed below, the tendency of some SBB-containing proteins to self-assemble enables this small module to elaborate new biological functionality.

### 1.2 SBBs as a unifying structural theme in many cellular pathways

The deep functional plasticity of SBBs seems to stem from several unique structural and physicochemical properties of this fold. Perhaps most important, small barrels are quite robustly ‘foldable’, and an unusually wide range of amino acid sequences can adopt this fold—i.e., a vast sequence space is compatible with this fold. A corollary of this principle is that a given β-barrel sequence can mutate significantly (e.g., evolutionary drift) without compromising the structural integrity of this fold; this property, in turn, makes the β-barrel fold a quite extensible platform for evolving, and potentially engineering, new functionalities. This evolutionary benefit comes at a practical price (for researchers): sequence similarity levels are often so low, even among SBBs of nearly identical backbone 3D structures, that identifying small barrel structures solely from sequence is frequently impossible (Theobald and Wuttke 2005; Dickey, Altschuler, and Wuttke 2013). Typically, new functional roles can be associated with specific structural features, but the SBB is more puzzling; in at least one SBB case (the OB fold), function is more closely correlated with sequence phylogeny than with the structural classification (Theobald and Wuttke 2005). Intriguingly, this phenomenon of a vast sequence and function space has been identified for some other folds involving β-sheets, including the eight-stranded TIM barrels (Nagano, Orengo, and Thornton 2002) and the β-sandwich framework of immunoglobulins (Bork, Holm, and Sander 1994).

SBBs from different functional classes are found in a variety of SCOP superfamilies, but common themes across these superfamilies—and, indeed, even potential relationships within a superfamily—have gone largely unreported. For instance, the Sm-like superfamily (b.38.1), which is strongly associated with mRNA splicing and processing, also contains (i) a domain from a membrane protein channel (family b.38.1.3), (ii) the hypothetical lipoprotein YgdR (b.38.1.6), and (iii) the bacterial RNA chaperone Hfq (b.38.1.2). Though in the same SCOP superfamily, it is difficult to conceive, for instance, of these three sets of proteins as being homologous. A profile-based phylogenetics approach has implicated divergent evolution as being at least partly responsible for the interrelationships among various small β-barrels (Theobald and Wuttke 2005); however, note that a failure to recognize potential evolutionary relationships, including possible convergent evolution towards the same fold, also could occur because information about SBB-containing proteins has been fragmented and disjoint in the literature. Such has been the case at least partly because of: (i) the remarkable functional diversity of these proteins; (ii) the small size and vast sequence space of this 3D fold, which hinders the detection of related (homologous) structures via sequence similarity beyond the twilight zone (Rost 1999); and (iii) SBBs have been assigned to many different SCOP folds. Thus, although SBB-containing proteins have been studied for many decades, someone studying, say, the Sm/LSm proteins in RNA splicing may be entirely unaware that these proteins adopt the same fold as the SH3 domain that binds polyproline-containing sequences in signal transduction cascades, the Chromo domain involved in chromatin remodeling, the ribosomal and other proteins involved in translation, or even the membrane channels involved in a bacterial cell’s response to a mechanical stress such as osmotic shock (e.g., the MscS protein).

To address this ‘knowledge gap’, here we synthesize and unify many lines of data, observations and analyses. In taking this first step towards developing a systematic and coherent model of structure↔function relationships among small β-barrel–containing protein superfamilies, we note that several distinct nomenclatures have arisen for describing the elements of the fold; these historical patterns reflect the isolation and lack of cross-talk between the various fields of study, as alluded to above. Also, note that classifying a newly identified small barrel structure can be baffling, from a structural bioinformatics perspective: sometimes a new structure is reported as similar to a specific—but not necessarily optimal—SCOP fold. Therefore, the present work largely consists of a thorough, survey-based analysis of the structural features of SBB domains. As part of our analysis, we systematize the various nomenclatures that have developed, we examine the anatomy of this structural fold, and we address how functional diversity can be achieved by such a seemingly simple structural unit (including via a multitude of oligomeric states and polymerization into fibrils). In places, we also highlight aspects of the SBBs that are hitherto unexamined, and which merit future investigation by experimental means. More broadly, this work helps systematically ‘define’ the unique SBB fold, and highlights the structural properties that enable its vast functional diversity; a detailed functional analysis of this fold will be presented elsewhere.

### 1.3 SBB cellular pathways: Evolution and functional diversity

The evolutionarily ancient SBB domain occurs in proteins from viruses, bacteria, archaea and eukaryotes, and it appears to act as a fundamental component in diverse biological pathways. SBB-based cell biology ranges from RNA biogenesis and decay/degradation pathways (splicing, RNAi, sRNA) (Wilusz and Wilusz 2013; Vogel and Luisi 2011; Mura et al. 2013; J.-B. Ma, Ye, and Patel 2004), to structural scaffolding and organization of chromatin DNA (Robinson et al. 1998), to maintenance of genomic integrity (Flynn and Zou 2010), to the translational apparatus (Klein, Moore, and Steitz 2004; Valle et al. 2002; Lomakin and Steitz 2013), and even to seemingly unrelated processes such as the host immune response (Cridland et al. 2012; Shaw and Liu 2014; Jin et al. 2012) and membrane transport. In addition to translation, splicing and other ancient/primitive pathways, SBB proteins also play key roles in pathways that likely evolved more recently (signaling and regulatory circuits, epigenetic modifications, etc.). As concrete examples, (i) the recognition of histone tails by SBBs underlies chromatin remodelling and the regulation of gene expression (Patel and Wang 2013); (ii) recognition of a polyproline signature motif makes the SH3 barrel a uniquely versatile adaptor/scaffold domain in regulatory cascades (McCarty 1998); (iii) eukaryotic Sm proteins form the common cores of small nuclear ribonucleoprotein complexes (snRNPs), and as such are key components of the spliceosome; and (iv) the bacterial Sm homolog, known for historical reasons as ‘Hfq’, is broadly involved in post-transcriptional regulatory pathways that hinge upon interactions between a small non-coding RNA (sRNA) and a target mRNA. This vast functional scope distinguishes SBBs from RRM domains, which also use a variety of molecular interaction strategies, but whose primary roles are associated with post-transcriptional steps in gene expression.

How early in evolution did the SBB fold arise, and does convergent evolution account for some instances of this fold in extant homologs (i.e., the fold arose multiple, independent times)? The SBB fold occurs in many ribosomal proteins and other ancient proteins involved in translation (Klein, Moore, and Steitz 2004; Valle et al. 2002; Lomakin and Steitz 2013), and it plays key functional roles in other basic (core) cellular pathways, such as the aforementioned roles in genome integrity and RNA processing. Thus, it is unsurprising that eliminating or compromising the functionality of many SBB-based proteins is associated with various cancers, inflammatory diseases, and other human ailments (e.g., the SBB-based Sm proteins were first discovered as the autoantigens in the autoimmune disease systemic lupus erythematosus (Tsokos 2006)).

SBB-containing proteins span a vast range of biochemical and cellular functionalities. How much of this versatility is directly attributable to the SBB domain itself? In analyzing structure↔function↔evolution relationships for an SBB domain embedded in a larger protein, one can try to delineate the specific molecular functionality of the SBB domain itself, as distinct from (yet contributing to) the net function(s) of the whole protein. The functionality of the SBB domain can be distilled into three overarching categories: (i) *Stabilizing macromolecular assemblies*, either by (a) serving as structural platforms/cores (examples include the roles of Sm and LSm oligomers in nucleating snRNP assembly, as well as verotoxin, HIN, TEBP and RPA proteins), or by (b) providing small stabilizing regions, as with individual SBB-containing ribosomal proteins, enmeshed in a network of rRNA structures. (ii) *Chaperoning RNA interactions* with other RNAs or proteins. Here, examples again come from the eukaryotic Sm/LSm proteins and, notably, the bacterial RNA chaperone, Hfq (Mura et al. 2013). Another notable example of facilitating RNA interactions is the role of Argonaute in binding to small, non-coding ‘guide’ RNAs in RNA silencing pathways (Gorski, Vogel, and Doudna 2017). Finally, SBB proteins also (iii) *Relay signals in biological pathways*, either by (a) being part of an adaptor or scaffold protein itself and thereby helping localize proteins to their target biological complexes (Good, Zalatan, and Lim 2011), such as for polyproline-binding by the SH3 domain and in the recognition of modified histone tails by the Chromo/Tudor domain, or by (b) providing allosteric regulation, in the case of ribosomal proteins S12 and Spt5 (Gregory, Carr, and Dahlberg 2009; W. Li, Giles, and Li 2014). Finally, there are also two reports of an enzyme’s catalytic residues lying within an SBB domain—namely, *E. coli* signal peptidase and the self-cleaving transcriptional repressor LexA (Luo et al. 2001; Paetzel, Dalbey, and Strynadka 2002).

### 1.4 Variability is the theme, there are no golden rules

A striking theme with the SBB domain is its great variability: In terms of sequence, structure, function and evolution, it seems that the only golden rule is that there are no golden rules (**GB Shaw**). As outlined with a few examples below, the SBB domain features extensive variability in terms of (i) ligand-binding properties and cellular pathways, (ii) 3D structures (variations on the fold), and (iii) oligomerization behavior and quaternary architectures.

The ligand-binding properties of SBB domains, either alone or as part of a multi-domain protein, exhibit great variation. SBB binding profiles for DNAs, RNAs, or other proteins vary from entirely generic (no sequence specificity), to partially nonspecific, to highly specific. An example of nonspecific binding is provided by the OB fold-containing <u>r</u>eplication <u>p</u>rotein <u>A</u> (RPA), which binds ssDNA and is involved in replication, recombination and repair (A. Bochkarev et al. 1997; Bochkareva et al. 2002). Examples of SBB-containing proteins with sequence-specific binding profiles include those that bind/protect telomere ends, e.g., the S. *cerevisiae* cell division control protein cdc13 (Mitton-Fry et al. 2004); the S. *pombe* POT1 (Lei, Podell, and Cech 2004), which binds a ^5′^GGTTAC^3′^ recognition site in ssDNA; and *Oxytricha nova* TEBP (Horvath et al. 1998). In each of these cases, the degree of sequence specificity, the sequence of the cognate recognition site (in cases of specific binding), and the binding affinity are dictated by the details of the an underlying structural mechanism which varies from one SBB-containing protein to another.

In terms of structural variability at the gross level of domain arrangement and general architecture, note that some SBB domains function as single, autonomous proteins, while in other cases the SBB module is part of a multi-domain protein. The former case is illustrated by several ribosomal constituents, such as the SH3-containing L14 protein, L21e, L24 (Klein, Moore, and Steitz 2004), and the OB-containing S12 and S17 (Brodersen et al. 2002). Most well-characterized Sm/LSm/Hfq proteins act as standalone domains, though bioinformatic studies of the domain architecture of Sm homologs suggest that C-terminal extensions, and even entire functional domains (e.g., putative methyltransferases), can be appended to some LSm homologs (Albrecht and Lengauer 2004). Many other SBB domains are also fused or embedded within larger proteins. As two examples, note that (i) many kinases contain an SH3 domain (Morton and Campbell 1994), and other proteins involved in signal-recognition contain Tudor and Chromo domains (Blus, Wiggins, and Khorasanizadeh 2011), while (ii) the RNA-binding ribosomal protein L2 consists of two fused SBB domains, with the N-terminal region adopting an OB-fold, while the C-terminal half is an SH3-like barrel (Diedrich et al. 2000). SBB-containing proteins also exhibit deep variability at the quaternary structural level. Many SBB proteins assemble into oligomers that act as the biological functional unit (e.g., toroidal Hfq discs that bind RNAs on either face). In some of these cases, the SBB module appears to enable oligomeric plasticity, between paralogs or even with the very same protein. As a striking example of this plasticity, the *Archaeoglobus fulgidus* SmAP2 protein forms both hexamers and heptamers, depending on solution-state conditions (a hexamer at low pH, without RNA, but a heptamer in the presence of U-rich RNA (Kilic et al. 2006)).

Intriguingly, some types of biological pathways are enriched in proteins containing SBBs. In these cases, the SBB functions in one of various modes, using different binding surfaces of the barrel and recognizing various binding partners (some bind to nucleic acids, others to proteins). A notable example is afforded by eukaryotic pre-mRNA processing pathways, where there appears to have been an evolutionary ‘fixation’ of the SBB fold in at least five distinct (functionally unrelated) steps along the intricate *snRNP assembly* → *spliceosome biogenesis* → *intron excision* pathway: (1) the canonical Sm hetero-heptamer scaffolds the formation of spliceosomal snRNP cores (U1, U2, etc.) (Salgado-Garrido et al. 1999); (2) a specific set of seven Sm-like (LSm) paralogs forms the hetero-heptameric U6 snRNP core (Salgado-Garrido et al. 1999), while other LSm heteromers form the U7 (histone-processing) and other RNP cores (Azzouz et al. 2005); (3) the Tudor domain is a key SBB found in the survival of motor neurons (SMN) protein, whence it interacts with methylated Arg side-chains in the tails of some Sm proteins in order to help assemble mature snRNP cores (Selenko et al. 2001); (4) the Gemin6/7 proteins also contain SBBs, and are key players in the eukaryotic snRNP biogenesis pathway (Y. Ma et al. 2005); and (5) the protein pICln, a methylosome subunit which is a 6-stranded SBB itself, chaperones a key step in the snRNP core biogenesis pathway (Grimm et al. 2013); intriguingly, the SBB module of pICln allows it to act via “molecular mimicry” of the SBB domain of the canonical/core Sm proteins (Grimm et al. 2013). Taken together, these features illustrate why the SBB domain is a rich and interesting system for studying the interrelationships between protein structure, function and evolution.

## 2. Results and Discussion

### 2.1 Scope of this work

Many protein folds are associated with the term ‘β-barrel’. For instance, 53 folds in SCOPe 2.06 (Chandonia, Fox, and Brenner 2017) are defined as a ‘barrel’ or ‘pseudo-barrel’, and 79 X-groups appear under the architecture of β-barrel in the ECOD classification (Cheng et al. 2014). The many SCOP (v.2.06) folds to which SBB proteins belong include the following (the number of associated superfamilies is given after the common name): (i) b.34 (SH3-like; 21 superfamilies), (ii) b.38 (Sm-like; 5 superfamilies), (iii) b.39 (ribosomal protein L14; 1 superfamily), (iv) b.40 (OB; 16 superfamilies), (v) b.136 (stringent starvation protein B; 1 superfamily), and (vi) b.137 (RNase P subunit p29; 1 superfamily). This selection of six SBB folds is biased towards those exhibiting pseudo-symmetry.

Barrels without internal symmetry, such as SCOP (v.2.06) folds b.35, b.36, b.41, b. 55, b.87 and b.138, are not treated here because of constraints on the length and scope of this work; notably, these are not highly populated folds, in terms of number of superfamilies, though some of them are abundant in the literature (e.g., b.35 is the GroES-like fold and b.36 is the PDZ domain) or exhibit intricate structural features (e.g., b.87, the LexA/signal peptidase domain, has an embedded SH3-like barrel). In terms of sequence diversity and functional breadth, the SBB can be technically termed a ‘superfold’ (Orengo, Jones, and Thornton 1994); in many ways, the SBB is comparable to another β-rich small superfold, namely ferredoxin (d.58, encompassing 59 superfamilies). How both the SBB and ferredoxin superfolds achieve such immense functional diversity (e.g., whether there are parallels between them) is an intriguing question for future work. Here, we focus on three SCOP folds—b.34 (SH3), b.38 (Sm) and b.40 (OB)—which represent the vast majority of known SBB structures and functions, and which offer insight into the structural and functional plasticity of small barrels.

### 2.2 General anatomy of small β-barrels

#### 2.2.1 Geometric and protein structural characteristics of the small β-barrel

In general, a β-barrel can be thought of as a β-sheet that twists and coils to form a closed structure in which the first and last strands are hydrogen-bonded (Murzin, Lesk, and Chothia 1994b, [a] 1994). Though they have not been formally (or precisely) defined in the literature, here we define **small β-barrels** (SBBs) as domains, typically ≈60-120-residues long, with a specifically superimposable framework of β-strands; often, the spatial pattern of strands exhibits two-fold rotational pseudo-symmetry, and the side-chains that emanate from this backbone scaffold give a structural ‘core’ of ≈35 residues. Classically, barrels are defined by the *number of strands*, **n**, and the *shear number*, **S** (McLachlan 1979; Murzin, Lesk, and Chothia 1994a). The magnitude of **S** describes the extent of stagger of the β-sheet or, equivalently, the tilt of the barrel with respect to its principal geometric axis; in turn, the magnitude of the stagger defines the degree of twist and coil of the strands, and is correlated with the internal diameter of the barrel (Murzin, Lesk, and Chothia 1994b, [a] 1994). An increase in the tilt of the barrel (i.e., **S**) is proposed to have occurred over the course of evolution (Caetano-Anollés and Caetano-Anollés 2003). Alternatively, barrels can also be viewed as two β-sheets packed face-to-face, with the strands in each sheet lying roughly perpendicular to one another (Chothia and Janin 1982). Such barrels have greater stagger values and are generally ‘flatter’ (cross-section through the barrel is more elliptical than circular), allowing the two opposite faces to pack closely together.

SBBs are of this more orthogonal barrel type, generally with few strands (low *n*) and high shear (*S* ≈ 2*n*). In SCOPe version 2.06, b.34 (SH3) and b.38 (Sm) are defined as *n*=4, S=8 with an SH3 topology, while b.40 (OB fold) is defined as n=5, S=10 (or S=8). In most cases the fourth strand, as defined in SCOP for b.34 and b.38, is interrupted by a short 3_10_ helix, resulting in two strands (e.g., the β4 and β5 strands of Hfq; Fig 1). Adhering to precedents that have been set by much of the literature, here we define small β-barrels as containing five strands, organized as two orthogonally-packed sheets. There are two distinct topologies of the small barrels treated here: SH3-like and OB-like sheets. As described in Figure 2, these two topologies are related via a (non-circular) permutation, resulting in the same 3D framework of strands. To lessen confusion, in what follows we use the SH3 nomenclature; the OB fold is treated in a separate section (‘Further structural variation’, Section 2.3.4 below).

**Figure 2.**
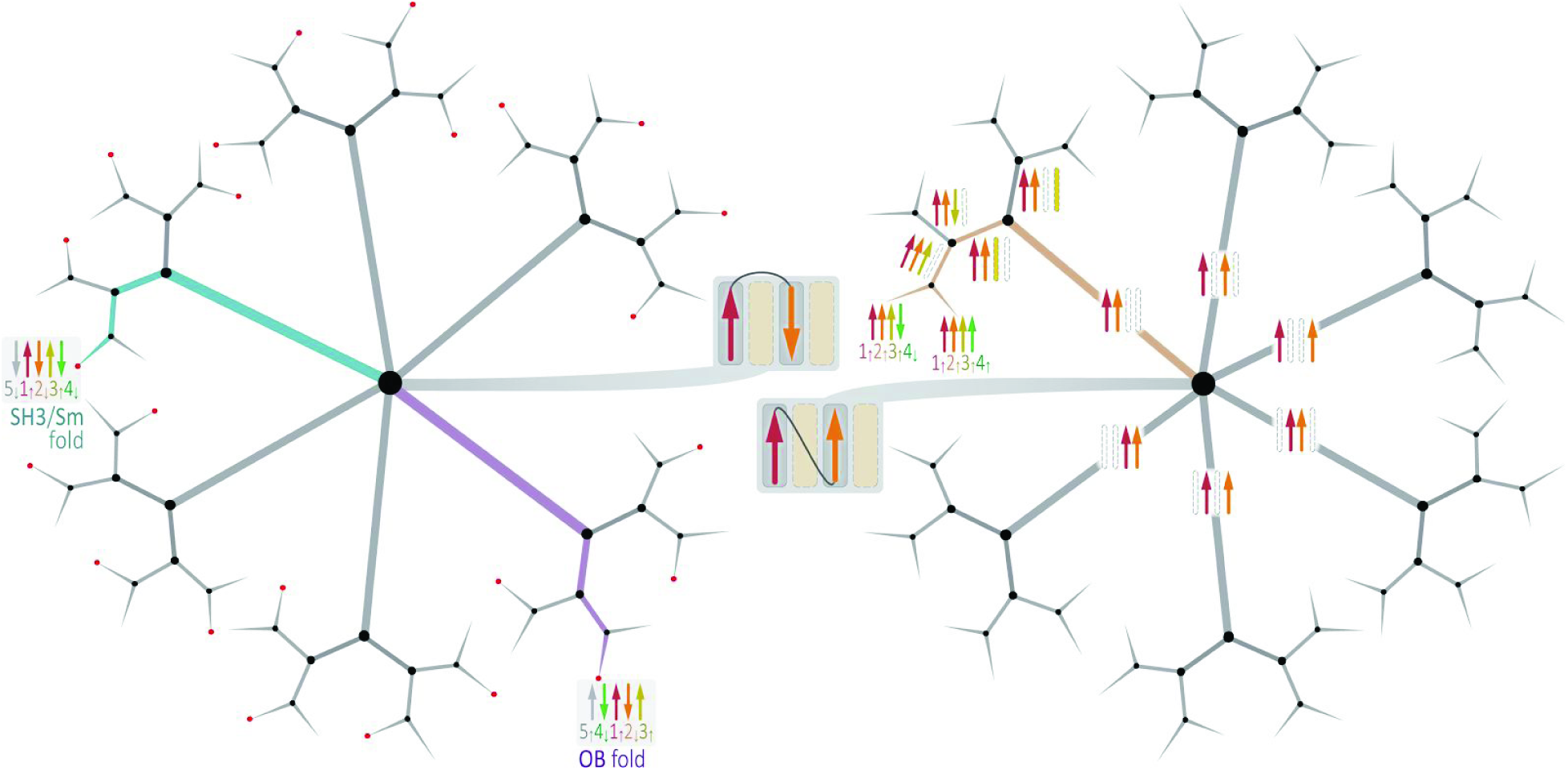
**Exhaustive enumeration of the 96 unique topologies of a four-stranded β-sheet,** via a decision tree-like approach, and the difference between the SH3 and OB domains. Here, strand β1 is red, β2 is orange, β3 is yellow, and β4 is green; for the SH3 and OB folds, the fifth strand is also shown (grey). Branches along a sample path in this digraph are highlighted in tan (subtree at right), yielding the 1↑2↑3↑4↑ and 1↑ 2↑ 3↑ 4↓, topologies (in Zhang & Kim’s nomenclature). The base of the overall tree (at the center) is a decision between the two possible configurations (parallel, antiparallel) for the simplest possible sheet—i.e., a tandem pair of strands (➜〰➜). Traversing the tree from this split ‘root’ to the leaves corresponds to building-up the sheet, and the tree’s branching structure elucidates the n!·^n−2^ unique topologies that are possible for a sheet of *n* strands; the successive branches of this unrooted *k*-ary tree are of degrees 2, 6, 2, 2, 2. The positions of the SH3/Sm and OB folds are indicated by cyan and purple paths (subtree at left). Other features of β-sheets are also elucidated by this hierarchical representation, such as the fact that there are 24 unique arrangements of two sequentially adjacent β-hairpin motifs (red circles, left subtree).

#### 2.2.2. Topological descriptions

A few structural features uniquely characterize the SH3-like SBB domain. The fold consists of five β-strands arranged in an antiparallel manner (Figs 1,2). A stringently-conserved Gly in the middle of strand β2 enables severe curvature of the backbone; this Gly is often followed by a β-bulge, dividing the strand into N-term (β2N) and C-term (β2C) segments (Fig 1A,B). As such, two orthogonal β-sheets (Sheet A, Sheet B) can be defined as comprising the β-barrel (Fig 1B); this view effectively makes the SBB a six-stranded barrel, with each β-sheet consisting of three β-strands. A short, single-turn 3_10_ helix links strands β4 and β5; the β4 and β5 strands straddle the barrel and belong to different β-sheets (as do β2N and β2C). This architectural arrangement enables multiple barrels to oligomerize via interactions of β4…β5’ of adjacent monomeric subunits (indicated by a prime)—a critical feature in forming toroidal discs (discussed further in this section).

The strands in an SBB are generally rather short, being ≈4-6 residues long (see below), and are connected by loops; the loops in some SBBs adopt known β-turn geometries (see, e.g., β-turn geometries mentioned in *A. aeolicus* Hfq (Stanek et al. 2017)), while others are more irregularly structured. Sheet A, also referred to as the *Meander* (Fig 1B), is a three-stranded β-sheet consisting of strands β2C, β3 and β4 (contiguous in sequence and in space). Sheet B, which consists of strands β5, β1 and β2N, and is non-contiguous (Fig 1B), links the C-terminal and N-terminal strands of the protein in an antiparallel fashion; for that reason, this sheet has also been referred to as the *N-C Sheet* (Fig 1B). Several other structural motifs and features have been described over the years for different classes of SBB domains, as summarized below.

*Proto-domains* (Fig 1C) are related by pseudo-symmetry within a single domain. They were noticed relatively early in the history of protein structure; for example, the six-stranded β-barrels of serine proteases were seen to exhibit C2 symmetry (McLachlan 1979). Some domains, such as those of serine or aspartyl proteases, are thought to have arisen from ancient duplications; in such cases, the sequence signal may be lost, while structural similarity persists and is more apparent. To our knowledge, proto-domains have not been described in small barrels, such as analyzed here. The SBB can be viewed as two proto-domains related by C2 symmetry. In the case of SBBs, proto-domain 1 consists of β1, β2N and β2C, while proto-domain 2 consists of β3, β4 and β5 (Fig 1C). Even if the existence of such proto-domains is merely a geometric byproduct of forming a closed barrel (via sheets like those shown in Fig 2), this 2-fold symmetry of the barrel does appear to be a recurring feature of SBB domains.

*The KOW motif* (Fig 1D) (Kyrpides, Woese, and Ouzounis 1996), which is found in some RNA-binding proteins (mostly small barrels in ribosomal proteins), consists of β1, β2, and the loops preceding β1 and following β2; together, this spans a total of 27 residues. A hallmark of this motif is alternating hydrophilic and hydrophobic residues with an invariant Gly at position 11 (Kyrpides, Woese, and Ouzounis 1996).

*Functional motifs* (Fig 1E) are exemplified by Sm-like proteins (b.38). Here, ‘function’ is meant generally, and in multiple senses—e.g., biochemical functionality (such as RNA-binding), or structural/physicochemical functionality (such as mediating interactions between subunits). The ‘Sm1 motif consists of β1→β3 and the ‘Sm2 motif consists of β4→β5, linked by a short, four-residue 3_10_ helix (Schumacher et al. 2002). The Sm2 substructure, with its β4-β5 strands straddling the barrel, is a significantly conserved feature, and possibly a signature of all small barrels with SH3-like topology. In fact, superimposing this pattern alone can yield high-quality structural alignments for the entire conserved structural framework (i.e., fold) of various SBB domains.

#### 2.2.3 The hydrophobic core and conserved structural framework

The hydrophobic core of the SBB is minimalistic, consisting only of the six elementary strands that form the conserved structural framework: β1, β2N + β2C, β3, β4 and β5 (Figs 1, 3). These strands are short, comprising roughly four to six alternating inside/outside residues, unless bulges are present. Only two strands, β1 and β3, are completely saturated in terms of their backbone hydrogen-bonding capacity. The structural framework of β-strands is the key property of SBB proteins and is well-captured by the Hfq barrel, where all loops are reduced to tight β-turns. The structural framework tolerates diverse residue replacement as long as a compact, well-packed hydrophobic core is preserved, as evidenced by interdigitated barrels, barrels inserted within each other, and barrels with deviation from the regular SBB fold (Section 2.5.1). Indeed, an antiparallel configuration of short β-strands may enjoy substantial ‘wiggle room’ for compensatory changes, while preserving overall structural integrity and thermodynamic stability—the register of some strands may shift, strand geometries may undergo minor rearrangements or adjustments, and so on.

**Figure 3.**
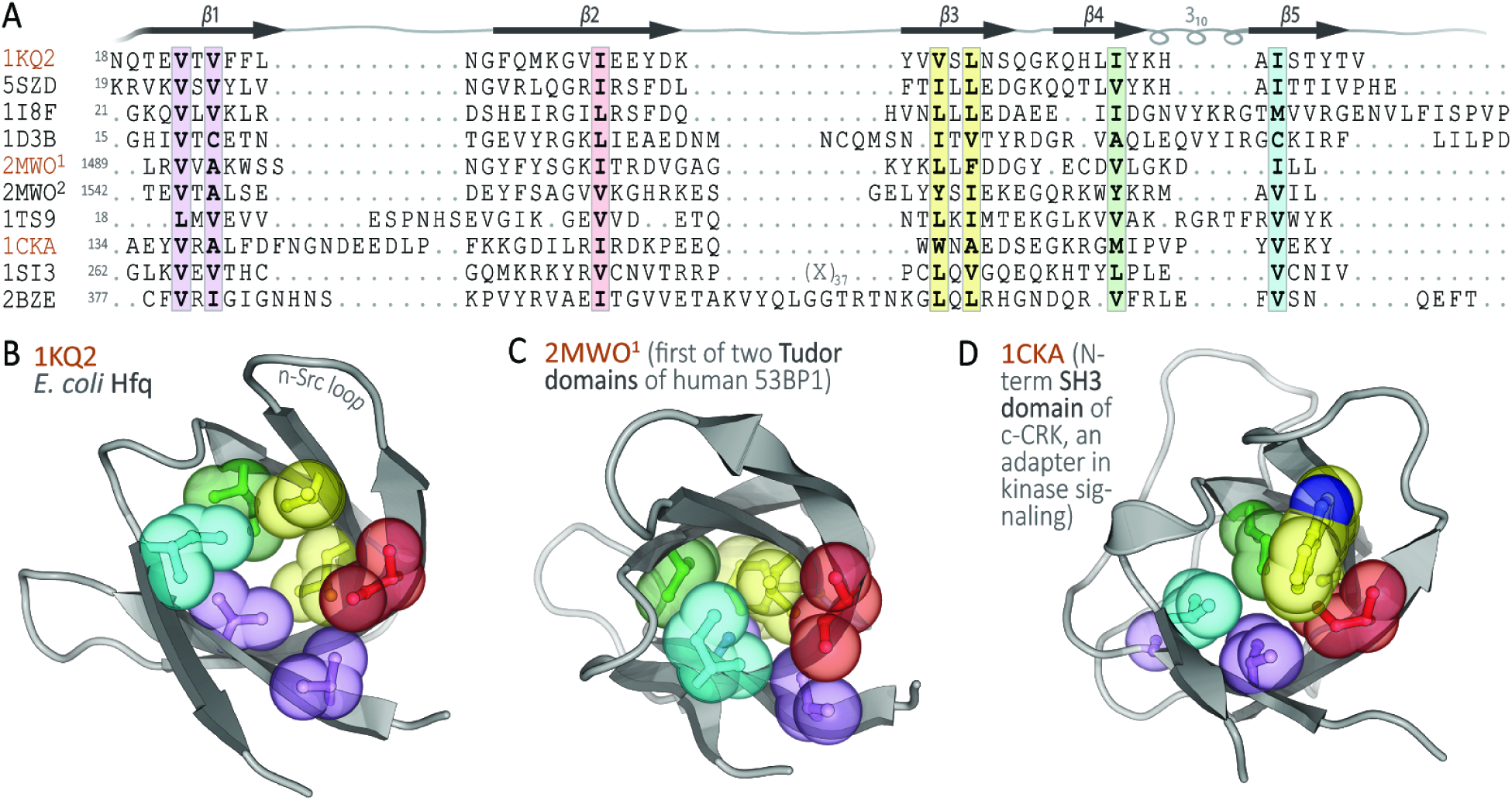
Hydrophobic core of the SBB. (A) A multiple sequence alignment of β-barrel homologs reveals several highly-conserved residues, highlighted here in color (Fig 4 shows the 3D structures of some of these proteins). At the sequence positions identified here, each β-strand contributes one or two conserved apolar residues to form the structural core of the barrel. For clarity, an unconserved 37-residue segment of 1SI3 is denoted ‘X_37_’, and superscripts distinguish the two Tudor domains that comprise the tandem repeats in 2MWO. The seven residues colored in this MSA are shown as space-filling spheres in the ribbon diagrams of (B) *E. coli* Hfq (1KQ2), (C) a Tudor domain (2MWO), and (D) an SH3 domain, as represented by 1CKA. In these panels, residues are color-matched to (A), two residues have the same color if on the same β-strand, and the β-barrel is drawn as a grey ribbon. The most conserved residues can be seen to form the barrel’s core, chiefly via hydrophobic packing interactions (London dispersion and other van der Waals forces).

Typically, each β-strand contributes one or two buried residues to the hydrophobic core (Fig 3). The two central strands—β1 at the center of Sheet B, β3 at the center of Sheet A (Fig 1B)—contribute two apolar residues each (yellow and magenta in Fig 3), while the four lateral strands (β2N, β2C, β4, β5) typically contribute one residue each to the hydrophobic core. Not all β2N strands (in all SBB structures) contribute consistently to the hydrophobic interior, so the minimal core can be taken as consisting of the seven residue positions shown in Fig 3. The conserved hydrophobic residue in β2C, typically a Val, Leu or Ile, follows Gly (the pivot point of the highly-curved β2 strand), and this residue is positioned at the beginning of the characteristic β-bulge. The hydrophobic residues in β4 and β5 abut the 3_10_ helix, either just before (β4) or after (β5) the helix. This hydrophobic core defines a stable, minimal SBB fold, leaving all solvent-exposed residues to interact with ligands or other biomolecules (and that, in turn, enables the diversification of function).

The seven-residue hydrophobic core can be extended in many ways (Table S1). Because the barrel is semi-open, various decorations can contribute hydrophobic residues to the minimalistic core. For example, the N-terminal helix in the Sm-like barrel extends β5-β2C of the otherwise open barrel. Similarly, the RT loop in the SH3-like barrel extends the β2N-β3 side of the barrel.

The outward-facing residues on the ‘edge’ stands—β2C and β4 in Sheet A (*Meander*), β5 and β2N in Sheet B (*N-C*)—can potentially form hydrogen bonds with other β-strands, unless they are sterically obstructed by terminal decorations or long loops. Such strand…strand interactions potentially have two effects. Firstly, they enable extension of the β-sheet of the barrel in the direction of the Sheet B face (Fig 1B), along loop L4 (in Sm terminology; or, the distal loop, in SH3 terminology); indeed, loop L4 is known to be highly variable in length and in sequence in SBBs of eukaryotic Sm and LSm homologs (see also the L4/distal loop of the SmD3 protein in Fig 4G). Secondly, such strand…strand interactions enable the formation of quaternary structures via hydrogen bonds and other atomic contacts, stitching together β4…β5 of adjacent subunits (see Sections 2.5.2 & 2.5.3, and Fig 8, covering oligomerization).

**Figure 4.**
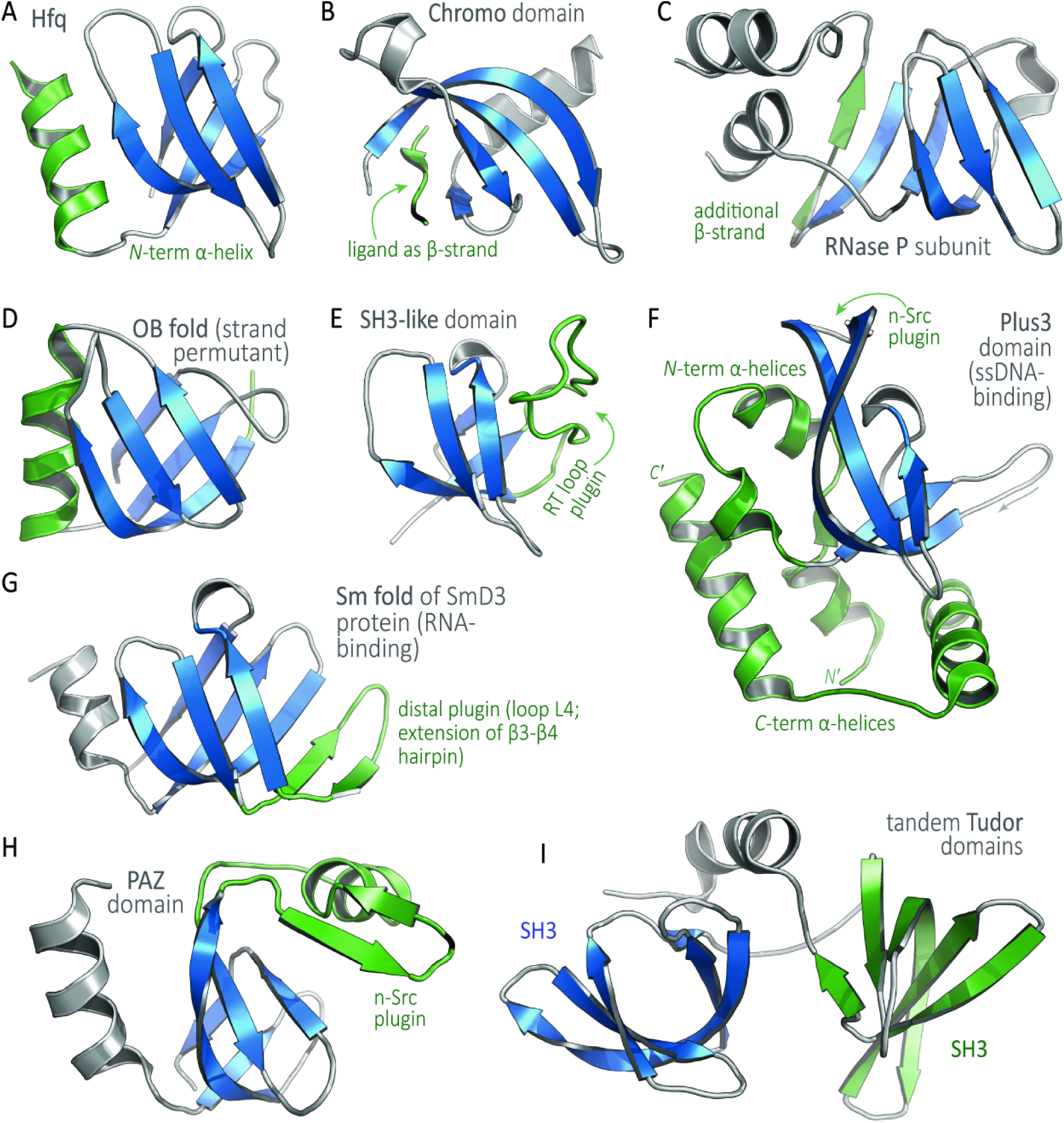
The SBB as a scaffold: Insertions, decorations, and other variations. In each panel, the SBB core is blue and variations are green. (A) The bacterial Sm protein Hfq (b.38.1.2) is taken as our reference structure (PDB 1KQ2), as it features the ‘cleanest’ and architecturally simplest barrel (e.g., minimal loops). (B) In this Chromo domain (b.34.13.1, PDB 1KNA), the SBB’s β5 strand is contributed by the cognate binding partner. (C) The RNaseP subunit P29 (b.137.1, PDB 1TS9) features an additional strand, β6, flanking the core SBB. (D) The nucleic acid-binding OB fold (b.40, PDB 1C4Q) is arranged as an SBB with a different sheet topology (see text and Fig 2). (E) An RT loop ‘plugin’ is found in the SH3 domain of the proto-oncogene c-Crk (b.34.2, PDB 1CKA). (F) In the DNA-binding Plus3 domain (b.34.21, PDB 2BZE), additional helices extend the N-and C-termini, and an n-Src plugin also occurs. (G) Many Sm folds feature a distal loop plugin, as illustrated here for the canonical, RNA-binding SmD3 protein (b.38.1.1, PDB 1D3B). (H) The PAZ domain, shown here from a human Argonaute protein (b.34.14, PDB 1SI3), has a plugin into the n-Src loop. (I) The Tudor-Tandem SH3 (b.34.9.1, PDB 2MWO) hastandemly repeated SBBs joined by a flexible linker; note that the relative spatial orientation of the two barrels in this single-chain structure can vary, in contrast to the rather precise geometric positioning of adjacent SBB subunits in Sm rings (Hfq hexamers, Sm heptamers, etc.).

##### 2.2.4 Hfq as a reference structure for all SBBs

Hfq, a key RNA-associated bacterial protein, adopts an Sm-like fold (SCOP b.38.1.2) and arguably represents the simplest example of an SBB (Fig 1 A). If one superimposes all small barrels and identifies structurally conserved regions (SCRs; Fig 3B), then Hfq appears to be the most regular structural representative (‘regular’ in the sense of clean and minimalistic—short loops, strands of roughly similar length, and minimal ‘decorations’ beyond the SBB’s obligatory five β-strands). Thus, for simplicity and clarity of presentation, we take Hfq as an archetypal representative of the SH3-like fold; note that Hfq is not assigned as such by SCOP, and also that other classification systems group many of the folds that share an SH3-like topology into a single category (Cheng et al. 2014). In short, Hfq provides a useful structural framework for all SH3-like folds, including the broad Sm superfamily.

#### 2.3 Beyond the SBB core: Loops, decorations, additional modules

##### 2.3.1 A brief overview and note on nomenclature

Unlike the core geometric framework of the small barrel—i.e., the all-β structural motif found in all the various SCOP folds that are SBB-like—all other structural elements, such as the loops, modules inserted within loops, and N’- and C’-terminal extensions, are highly variable (Fig 4). That the SBB fold can tolerate such variation is critical to its biological roles: these additional structural elements largely delineate the specific cellular functions of different SBB-containing proteins, irrespective of whether the structural similarity between these proteins stems from divergent evolution (i.e., homology) or, alternatively, convergent evolution.

Before analyzing the loops of the small β-barrels, it is worth noting that several different systems of nomenclature have arisen in research communities working on different (in terms of cellular functions) subsets of the universe of all small β-barrels. While the terminology for the β-strands is consistent (β1, β2,…), several naming schemes have emerged for the loops, thus muddying efforts at comparative analyses of different functional classes. The three most prominent nomenclatures are outlined in Table 1. First are the SH3-like barrels involved in signal transduction through binding to polyPro motifs (b.34.2), as well as chromatin remodeling via recognition of specific modifications on histone tails by Chromo-like (b.34.13) and Tudor-like (b.34.9) domains. Second are the Sm-like barrels involved broadly in RNA processing (b.38.1). Third are the OB-fold barrels (b.40), primarily involved in maintenance of genome integrity via binding to nucleic acids and oligosaccharides. To be consistent throughout this paper as well as inclusive of prior work, we cross-reference these terminological systems in Table 1, and we use the nomenclature for the SH3-like fold throughout, for either SH3-like (b.34) or Sm-like (b.38 or any other small barrel sharing the same topology, e.g. b.136, b.137 and b.39). Given the large volume of existing literature on OB proteins, its nomenclature is preserved here (with mapping to the SH3-like fold, when appropriate).

**Table 1.** Mapping the names of SBB loops, as used in the major superfamilies that share an SH3 topology, onto the SH3-like (b.34) notation used in this work. The SH3/Sm topology, using SH3 domain terminology, runs (α1–β1)–(β2–β3–β4)–β5, where β2–β3–β4 is a meander (see also Fig 2). A complete description of OB-fold loops is given in Table 2.

| Strands bracketing the loops (using SH3-like numbering) | **SH3-like** (b.34)<br>loop name | **Sm-like** (b.38)<br>name of<br>corresponding loop | **OB-fold** (b.40)<br>name of<br>corresponding loop |
| --- | --- | --- | --- |
| α-helix$\rightarrow\beta$1, or β0$\rightarrow\beta$1 | N-term loop | L1 | L01 |
| β1$\rightarrow\beta$2 | RT | L2 | — |
| β2$\rightarrow\beta$3 | n-Src | L3 | L12 |
| β3$\rightarrow\beta$4 | Distal | L4 | L23 |
| β4$\rightarrow\beta$5 | $3_{10}$ helix | L5 | — |

##### 2.3.2 Specific loop variations, including insertion of secondary structural elements

Loops that connect the β-strands in SBBs vary significantly in length and confer a plethora of functional roles (see below). SBBs consist of five, or sometimes six, loops. The first loop precedes the first strand, β1. The central four loops are always present (Fig 1): RT, n-Src, Distal, and 3_10_ helix (as defined for the SH3-like fold, b.34.2). Of these four loops, significant variations in the lengths of three—RT, n-Src and Distal—have been observed, and can be linked to specific biochemical functions. The fourth loop is almost always a short (single-turn) 3_10_ helix and is, infrequently, either (i) a distorted helix, as in RPP29 (Sidote, Heideker, and Hoffman 2004) or (ii) replaced by a longer loop, as in the TrmB proteins (Krug et al. 2006). Elongated loops often house additional secondary structures, as described below. A sixth loop is also possible (the C-terminal loop), linking the last β-strand to a C-terminal extension (e.g., a mixed α/β domain in *P. aerophilum* SmAP3 (Mura, Phillips, et al. 2003)) or to a sixth strand of the barrel, if present, as in the RNase P subunit shown in Fig 4C.

<u>RT loop, linking β1→β2</u> (Fig 4E): Long inserts into the RT loop, which links strands β1 and β2, results in the classical SH3 domain (b.34.2) that is ubiquitous in signal transduction. The SH3 domain binds proline-rich sequences using the elongated RT loop (as well as the n-Src loop and 3_10_ helix). The RT loop lies along the side of the barrel and caps one of its ends (Lim 1996); (Yu et al. 1992). Different pairs of loops form various pockets. In the PAZ domain (b.34.14) of the Piwi and Argonaute (RNA interference) proteins, aromatic residues of the elongated RT loop (Fig 4H) are part of an aromatic pocket formed between the loop and the α/β module (inserted into the n-Src loop, see below); this pocket laterally secures the RNA substrate (J.-B. Ma, Ye, and Patel 2004).

<u>n-Src loop, linking β2→β3</u> (Fig 4F, H): An elongated n-Src loop is observed in two functional families. The first case is that of the PAZ domains (b.34.14) of Piwi and Argonaute (J.-B. Ma, Ye, and Patel 2004), described in the previous section. In the case of the Plus3 domain (b.34.21) of the transcriptional elongation factor Rtf1, the extended n-Src loop contains two short (three-residue) β-strands and is involved in binding single-stranded DNA (de Jong et al. 2008).

<u>Distal loop, linking β3→β4</u> (Fig 4G): Perhaps the most notable example of elongation of the distal loop is with the eukaryotic Sm proteins, which are a core part of the splicing machinery. Elongation of this loop corresponds to extension of the β-hairpin formed by strands β3 and β4. In extending the distal loop, these two long β-strands become bent, similarly to β2, and can be seen as β3N and β3C, β4N and β4C (Kambach et al. 1999). Like β2, they simultaneously contribute to the formation of two sheets (Fig 1E). This extended β3-β4 hairpin (Fig 4G) results in a much larger hydrogen-bonded Sheet B, now containing the five strands β5, β1, β2N, β3C and β4N. The original Sheet A remains the same. Much of the sequence variation among Sm proteins occurs within the distal loop, and as extensions of it (see, e.g., the SmB protein and its alternatively-spliced variants).

<u>The 3_10_ helix, linking β4→β5</u>: The 3_10_ helix that connects strands β4 and β5 is generally short (*≈4* residues) and relatively invariant in structure. The geometry of this linker defines the relative positions of strands β4 and β5 which, in turn, generally straddle the barrel (the ‘edge strands’). Indeed, the dynamical flexibility and plasticity of this element is limited by the structural constraint that it link the β4〰β5 strands. This 3_10_ helix is present in virtually all SH3-like folds, but is absent in the OB fold for topological reasons (see below). In Sm proteins, the occurrence of this linker helix does not adhere to a strict pattern—it is absent in many eukaryotic Sm structures and Sm-like archaeal protein (SmAP) homologs, but present in most bacterial Sm (Hfq) protein structures. Intriguingly, sac7d, sso7d and other histone-like small archaeal proteins feature a second 3_10_ helix in the middle of the highly-bent β2 strand (Robinson et al. 1998), in place of the stereochemically-forgiving Gly that is phylogenetically conserved at that position in the Sm fold (Mura et al. 2013) and in many other SBB-containing proteins.

##### 2.3.3 N- and C-terminal decorations, capping of the barrel, small internal modules

Helices and additional loops or extensions often occur at the N-and C-termini in SBBs, and these are sometimes termed ‘decorations’. Their position relative to the barrel core varies. In some cases they affect the ability of the barrel to oligomerize. These decorations, in a manner similar to loop insertions, almost always have a functionally significant role. The following select examples illustrate how SBB decorations can serve as functional adaptations.

<u>N-terminal α-helix</u> (Fig 4A): The Sm-like fold (b.38) generally features an N-term helix that links to the main body of the barrel via a short loop. This region can engage in multiple interactions with both RNAs and proteins. The helix stacks against the open barrel and, in the context of an intact, hexameric Hfq toroidal disc (Sauer 2013), it lies atop the so-called *proximal face* (this face of the disc is defined below in Section 2.5.3; it has been termed the Loop L3 face for SmAPs (Mura et al. 2013)). In Hfq (b.38.1.2), the SBB’s α-helix mediates interactions with cognate sRNA molecules via a patch of conserved basic residues (Arg16, Arg17, Arg19) and a Gln8 ((Schumacher et al. 2002; Stanek et al. 2017); note that *E. coli* Hfq residue numbering is used here). A similar mode of RNA-binding appears to be conserved in the Sm-like archaeal proteins (Mura, Kozhukhovsky, et al. 2003; Thore et al. 2003). In LSm proteins, the N-term α-helix interacts with proteins Pat1C in the LSm1→7 (D. Wu et al. 2014) ring, and with prp24 in the LSm2→8 ring (Karaduman et al. 2008). In the case of Sm proteins (b.38.1.1), the same α-helix interacts with the β-sheet of the adjacent protomers during ring assembly (Kambach et al. 1999). For the eukaryotic paralog SmD2, a long N-terminal region harbors an additional helix (h0) that interacts with U1 snRNA as it traverses into the lumen of the heptameric Sm ring (Pomeranz Krummel et al. 2009; J. Li et al. 2016).

<u>C-term α-helices</u> (Figs 4C, F) can either augment existing binding interactions or mediate contacts with additional binding partners. For the LSm1→7 ring (b.38.1.1), a long helix formed by the C-term tail of the LSm1 subunit lies across the central pore on one face of the ring, preventing the 3’-end of RNA from exiting via that distal surface (Weichenrieder 2014). Notably, the novel structure of an Sm-like pentamer of putative cyanophage origin (Das et al. 2009) revealed that this homolog lacks an N-terminal helix, and instead features a well-defined C-terminal helix.

<u>N-term and C-term α-helices together</u> (Figs 4C, F) can interact to form a supporting structure/subdomain around the barrel, as in the case of the Plus3 (b.34.21) domain of Rtf1 (de Jong et al. 2008); in that system, three N-term α-helices and a C-term α-helix form a four-helical cluster that packs against one side of the barrel. The role of these helices is unclear, but the conservation of many residues in that region implies some presumptive functional significance.

<u>C-term tails</u> have, among all conceivable decorations, the least stereochemical and overall structural constraints. These regions can remain disordered and can vary significantly in length—for instance, >40 residues in SmD1 and SmD3, and >150 residues in SmB/B’ (Kambach et al. 1999). In the case of Sm proteins (b.38.1.1), the C-terminal tails of SmB/B’, SmD1 and SmD3 harbor RG-rich repeats that are critical for assembly of the Sm SBBs into a toroidal disc; assembly occurs via an intricately-chaperoned, arginine-methylation-dependent biogenesis pathway (Selenko et al. 2001; Friesen et al. 2001; Grimm et al. 2013). In Hfq (b.38.1.2), the disordered C-term tails are proposed to extend outward from the ring and mediate contacts with various RNAs (Beich-Frandsen et al. 2011), perhaps as an instance of a ‘fly-casting’ mechanism between a disordered region and its cognate ligand (Shoemaker, Portman, and Wolynes 2000; Levy, Onuchic, and Wolynes 2007). Most recently, it has been demonstrated that acidic C-terminal tails of *E. coli* Hfq interact with residues of the SBB core domain. This enables auto-regulation of the annealing between sRNA and mRNA by assisting the release of sRNA●mRNA pairs, increasing specificity of sRNA binding and preventing dsDNA aggregation on the rings (Santiago-Frangos et al. 2017); the latter property is important, as at least some fraction of Hfq, which exists at high intracellular concentrations, is thought to colocalize with the bacterial nucleoid. Finally, we note that in some (underinvestigated) LSm homologs, lengthy regions—of up to hundreds of residues—extend the C-termini well beyond the SBB core. At least five novel groups of homologs (LSm12→16) were bioinformatically detected in eukaryotes (Albrecht and Lengauer 2004); these extended SBBs likely act in RNA metabolic pathways (mRNA degradation, tRNA splicing, etc.), and the C-terminal regions in some of them have been identified as encoding putative methyltransferase activities. In these extended LSm homologs, the SBB acts as a module that imparts a specific functionality (e.g., nucleic acid-binding).

<u>Small internal modules</u> (Figs 4F, H) are short secondary or super-secondary structures (α/β or purely α) inserted within the loops of an SBB. These structural elements typically form a pocket against the barrel and are an integral part of barrel function. Examples include an α-β-β module inserted into the n-Src loop of the PAZ domain (b.34.14) (J.-B. Ma, Ye, and Patel 2004) and a β-hairpin extension module in the n-Src loop of the Plus3 domain (b.34.21) of Rtf1 (de Jong et al. 2008). Insertions can be entire, domain-sized modules (see Section 2.5.1.4)

##### 2.3.4 Further structural variations

The following vignettes briefly describe further structural variations that have been discovered by determining structures of SBB-containing proteins, chiefly via X-ray crystallography or NMR spectroscopy.

<u>Additional β-strands</u>: The example of RNase P subunit Rpp29 (Fig 4C; b.137.1) illustrates that the barrel core can be extended by a sixth strand; here, an extra β-turn in the C-terminal region (between strands β5-β6) is followed by the sixth β-strand, thus extending Sheet B (N-C) to four antiparallel strands (β6, β5, β1, β2N; (Sidote, Heideker, and Hoffman 2004; Numata et al. 2004)).

<u>Missing β-strands</u>: In at least one case (Fig 4B), namely the Chromo domain HP1 (b.34.13.2), the intact SBB is formed only upon binding of the cognate peptide ligand. HP1 exists as a three-stranded sheet A (meander), and the β-strand conformation of the peptide ligand templates the formation of the second β-sheet (N-C), thereby completing the barrel (Jacobs and Khorasanizadeh 2002).

<u>OB fold (b.40; Figs 4D, 5)</u>: An example of similar architectures, but differing topologies. Similar to the SH3-like barrel, the OB fold is a barrel comprised of a five-stranded antiparallel β-sheet. However, the SH3 and OB sheet topologies differ (Figs 2, 5): The SH3-like and OB topologies are related by a (non-circular) permutation, as noted previously (Agrawal and Kishan 2001; Theobald and Wuttke 2005). Our reference Sm-like fold, i.e. the Hfq protein, is well-suited to comparisons with the OB fold, as both topologies (Fig 2) are evidently compatible with the same 3D structural framework of β-strands (Fig 5). To avoid confusion between OB and Sm/SH3, we use OB strand mapping when discussing OB folds; that mapping is given in Table 2A.

**Figure 5.**
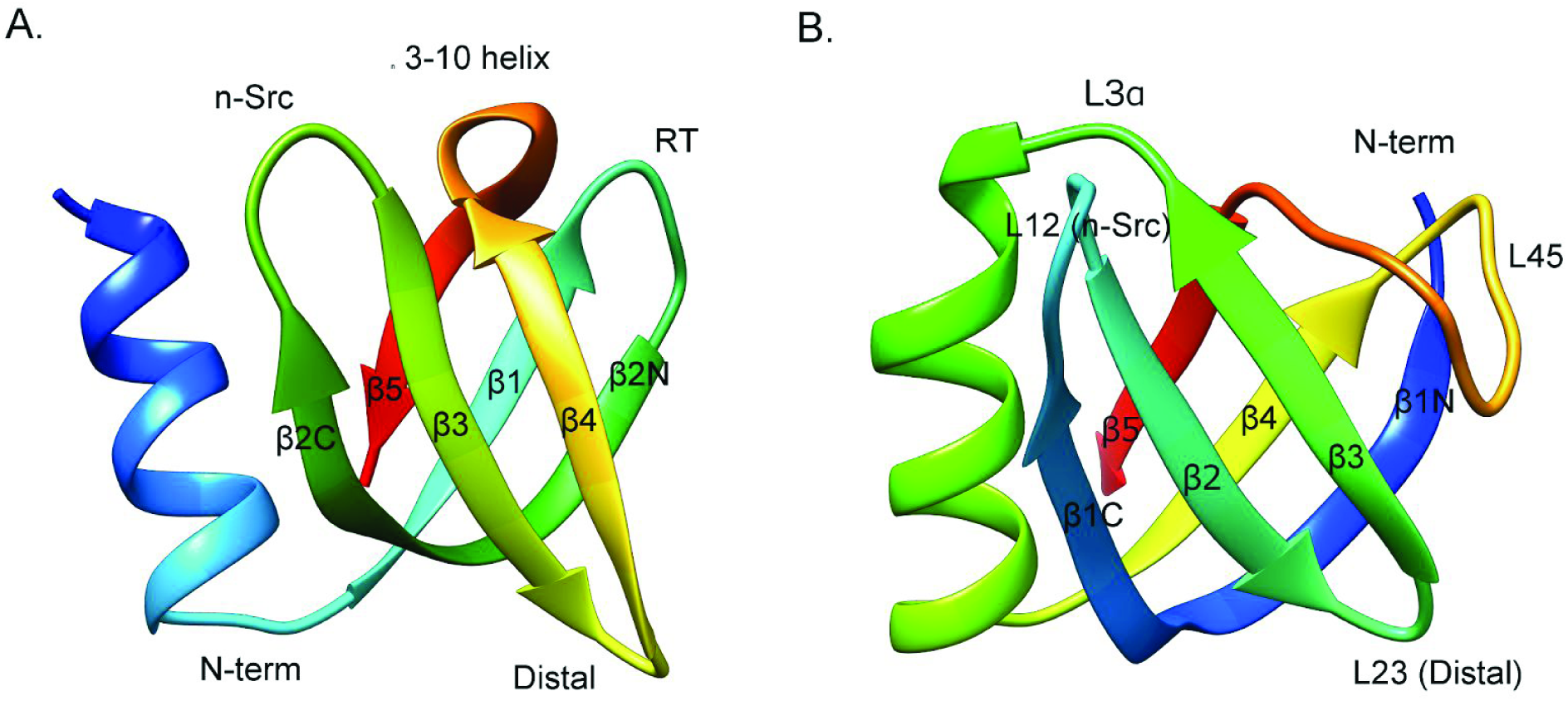
**Comparison of the OB and SH3-like folds,** with a mapping of strands and loops. In these ribbon diagrams, strands are labelled sequentially and colors progress from blue (N-terminus) to red (C-terminus). The SH3-like fold of Hfq (b.38.1.2) is shown in (A), and panel (B) illustrates the OB fold.

The correspondence between the OB and Sm-like folds is particularly striking, and notably both of these folds lack an α-helix found in the SH3-like fold. The permutation from the Sm-like fold to the OB-fold (Fig 2) places the N-terminal α-helix and β1 of Sm after the *Meander* [β2C-β3-β4] and before β5; thus, the initial Sm-like topology, [α1-β1]-[β2N-β2C-β3-β4]-[β5], results in the final OB topology [β2N-β2C-β3-β4]-[α1-β1]-[β5] (to use the Sm numbering throughout). Renumbering the permuted strands, using OB-fold nomenclature, then gives [β1N-β1C-β2-β3]-[α1-β4]-[β5]. The non-circular permutation preserves *Sheet A* (the *Meander*) in both topologies: [β2C-β3-β4] in Sm-like/SH3-like, and [β1C-β2-β3] in the OB-fold. Structural alignment of [β2N-β2C-β3-β4-β5] in SH3-like and [β1 N-β1 C-β2-β3]+β5 in OB yields an RMSD of 1.37 Å, using Hfq (PDB 1KQ1) as an SH3-like fold and verotoxin (PDB 1C4Q) as an OB-fold representative (and neglecting strand β1 of the SH3 fold and the analogous strand, β4, from the OB).

Of the five loops in the OB-fold (Table 2B), L12 can be clearly structurally mapped onto the n-Src loop and L23 to the Distal loop of the SH3-like fold (Fig 5). There is no good structural correspondence between the other loops. The RT and 3_10_ helix are absent from the OB fold but present in SH3-like topologies; conversely, L3α, Lα4 and L45 are unique to the OB fold.

**Table 2.** Mapping corresponding β-strands (A) and loops (B) between SH3-like and OB folds. The OB topology, using OB domain nomenclature, runs(β1-β2-β3)-(α-β4)-β5 where β2-β3-β4 is a meander.

| OB-fold strands | SH3-like fold strands |
| --- | --- |
| $\beta 1$ | $\beta 2$ |
| $\beta 2$ | $\beta 3$ |
| $\beta 3$ | $\beta 4$ |
| $\beta 4$ | $\beta 1$ |
| $\beta 5$ | $\beta 5$ |

**Table 2.** Mapping corresponding β-strands (A) and loops (B) between SH3-like and OB folds. The OB topology, using OB domain nomenclature, runs( $\beta 1$ - $\beta 2$ - $\beta 3$ )-( $\alpha$ - $\beta 4$ )- $\beta 5$ where $\beta 2$ - $\beta 3$ - $\beta 4$ is a meander.
| OB-fold strands | OB loops | Equivalent SH3 loops |
| --- | --- | --- |
| $\beta 1$-$\beta 2$ | L12 | n-Src |
| $\beta 2$-$\beta 3$ | L23 | Distal |
| $\beta 3$-α | L3α | — |
| α-$\beta 4$ | L$\alpha 4$ | — |
| $\beta 4$-$\beta 5$ | L45 | — |

The hydrophobic core of the OB fold contains the same seven residue positions defined for the SH3-like fold (Fig 3B, C, D), but is typically larger by virtue of (i) strand elongation (especially those bounding L12/n-Src), and (ii) formation of a possible hairpin within L45, which would then extend *Sheet A* by two β-strands. Most of the notable loop variations in the OB-fold are similar to those of the SH3-like fold, as summarized in the following vignettes.

<u>Insertion into the n-Src loop</u>: In the OB2 of BRCA2, a Tower domain is inserted in the L12 loop (corresponding to the n-Src loop). The Tower domain, which has been implicated in DNA-binding, is a 154-residue long insert that consists of two long α-helices and a three-helix bundle positioned between them (Yang et al. 2002; Alexey Bochkarev and Bochkareva 2004). In the C-terminal DNA-binding domain (DBD-C) of RPA70, a zinc-finger motif consisting of three short β-strands is inserted into the L12 loop of this OB fold (Bochkareva et al. 2002).

<u>Extension of the Distal loop</u>: The DNA-binding OB domain of cdc13 contains a unique pretzel-shaped loop L23 (corresponding to the Distal loop) that significantly extends the potential interactions of this barrel with DNA. This 30-residue–long loop twists and packs across the side of the barrel, and interacts with the L45 loop (Mitton-Fry et al. 2004).

<u>Change of internal α-helix and replacement with an Ω-loop</u>: In the DBD-C of RPA70, the α-helix lying between β3 and β4 is replaced by a helix-turn-helix, while in the DBD-D of RPA32 the same α-helix is missing altogether and is replaced by a flexible Ω-loop (Bochkareva et al. 2002).

### 2.4 Sequence variation, and electrostatic properties of SBB surfaces

In addition to variations in structure, variations in sequence further distinguish different barrels. Small barrels are extremely tolerant to mutations (see Section 2.6 on folding and stability of SBBs), and a common evolutionary strategy appears to have been the modulation of electrostatic interactions by changing the residues exposed in loops, sheets and decorations. In some cases, this means a shift in various physicochemical properties of residues—acidic/basic (positively/negatively-charged), polar/apolar, bulky/compact, etc. Such sequence changes can alter the properties of protein surface patches, or even entire sheets, yielding drastically different ligand-binding profiles, DNA/RNA interactions, and downstream physiological functions.

An interesting case is that of the HIN domains of AIM2 and p202, which bind dsDNA and are involved in the innate immune response (Fig 6) (Shaw and Liu 2014)(Jin et al. 2012; Q. Yin et al. 2013). Each HIN subunit consists of tandem OB-fold barrels, which are known to bind single-stranded, double-stranded, and quadruplex DNA with various affinities. Even though there is 36% sequence identity between the HIN domains in AIM2 and p202, the DNA-binding modes are entirely different (Fig 6A), largely because of variations in the electrostatic charge distributions. In the case of AIM2, the binding is mediated by positive charges on the convex surface of the barrel (Fig 6B). In the case of p202, the analogous surface is negatively charged and therefore does not interact with DNA. Instead, DNA is bound by the positively-charged loops (of the second OB barrel), on the opposite side of the barrel (Fig 6C). The same loops bear hydrophobic residues in the case of AIM2, and thus do not bind DNA (Jin et al. 2012; Q. Yin et al. 2013). The different binding surfaces correspond to different DNA-binding affinities, enabling these two proteins to act in a physiologically antagonistic manner.

**Figure 6.**
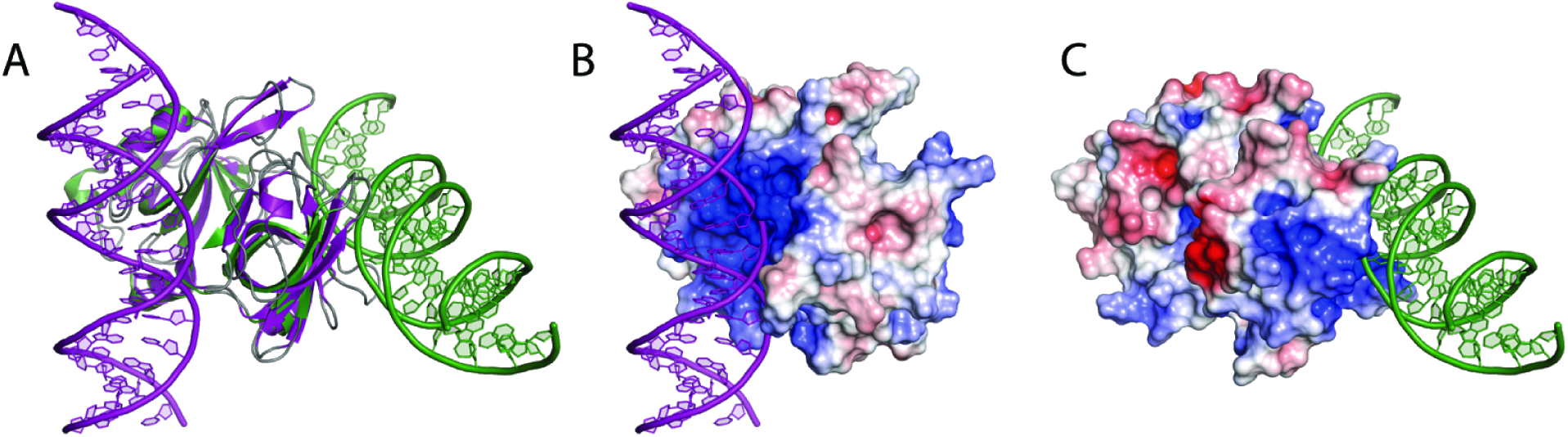
Different electrostatic charges on the surfaces of two very similar HIN domains. The HIN domain consist of two tandem OB-fold barrels. (A) Superimposed structures of the HIN domains of AIM2 (purple, PDB ID 3RN5) and of p202 (green, PDB ID 4LNQ) bound to double-stranded DNA (green double helix interacts with p202, purple double helix interacts with AIM2) via two distinct and separate interfaces. (B) Same view as in A showing only AIM2. The HIN domain of AIM2 is now shown as a surface representation and colored according to electrostatic potential. DNA binds to the first OB-fold barrel of the HIN domain; the interacting surface is positively charged (blue). (C) Same view as in (A), but showing only p202. The HIN domain of p202 is now shown as a surface representation and colored according to electrostatic potential. DNA binds instead to the positively charged patch (blue) on the second OB-fold barrel of p202.

Perhaps the most well-studied illustration of electrostatic variations in a small barrel concerns the charge distribution within the RT loop of polyproline-binding SH3 domains. Acidic residues in the RT loop, along with basic residues in the polyproline signal, mediate key electrostatic interactions; in addition, hydrophobic interactions involve the prolines themselves. The position of the basic residue (Arg) determines the orientation of polyPro peptide binding (Lim, Richards, and Fox 1994). The binding strength can be modulated by the number of acidic residues and their positioning within the RT loop (X. Wu et al. 1995). In at least one case—namely, interaction of the SH3 domain of Nck with the polyproline tail of CD3ε—binding can be switched on/off by simply phosphorylating a key tyrosine in CD3ε. This post-translational modification results in electrostatic repulsion between the poly-Pro region of CD3ε and acidic residues in the RT loop of the SH3 domain of Nck (Takeuchi et al. 2008), thus diminishing binding.

### 2.5 Joining barrels, covalently (in tandem) and noncovalently (as oligomers)

Small barrels tend to associate with one another at different structural scales. Interactions between tandem barrels within a single polypeptide chain are common, especially in RNA-binding proteins (Lunde, Moore, and Varani 2007; Cléry, Blatter, and Allain 2008) and in proteins that act as scaffolds for the binding of other proteins or nucleic acids (Good, Zalatan, and Lim 2011). Many proteins that consist of only an SBB domain are known to assemble into multimeric rings that function in many RNA-associated pathways, across all three domains of life. Though not ubiquitous among SBBs, the property of oligomerizing into rings is rather common for small barrels involved in RNA biogenesis (e.g., pre-mRNA splicing), as well as other RNA-associated pathways (e.g., the tryptophan-activated RNA binding attenuation protein, TRAP; reviewed in (Lunde, Moore, and Varani 2007)). Finally, small barrels (and oligomers thereof) have been found to self-associate into closed higher-order assemblies (e.g., head-head stacks of rings, with dihedral symmetry) or, in some instances, open-ended polymeric fibrils (e.g., head-tail stacking of rings of SBBs). Large supramolecular assemblies of SBBs are described in more detail below (Section 2.5.4).

#### 2.5.1 Tandem, embedded and enmeshed barrels

This section treats several combinations of β-barrels that occur either in tandem or are intertwined, within one protein chain.

##### 2.5.1.1 SH3〰SH3: Tandem Tudors

SH3-like barrels that are repeated in tandem often form barrel-to-barrel interfaces, and these can be constructed in various ways. Different linkers and sequences can lead to a number of tandem interfaces with varying extended sheets, allowing great plasticity. For example, in p53-binding protein 1 (53BP1, Fig 7A), hydrogen-bonding between β2N of the first barrel and β5 of the second one joins individual three-stranded β-sheets into an extended six-stranded sheet. The C-terminal α-helix strengthens the connection by interacting with multiple β-strands of both barrels (Charier et al. 2004). The tandem Tudor-like Agenet domains of FMRP associate with one another via interactions between each domain’s β2N. In the transcription elongation factor Spt5, which has five tandemly repeated KOW-containing Tudor domains, interactions between Tudor-2 and Tudor-3, which move as a single body, occur through β5 of Tudor-2 and residues immediately following β5 in Tudor-3 (Meyer et al. 2015). In the DNA/RNA repair protein KIN17, this interface is formed by N-terminal and C-terminal tails that interact with the linker between the two barrels (le Maire et al. 2006).

**Figure 7.**
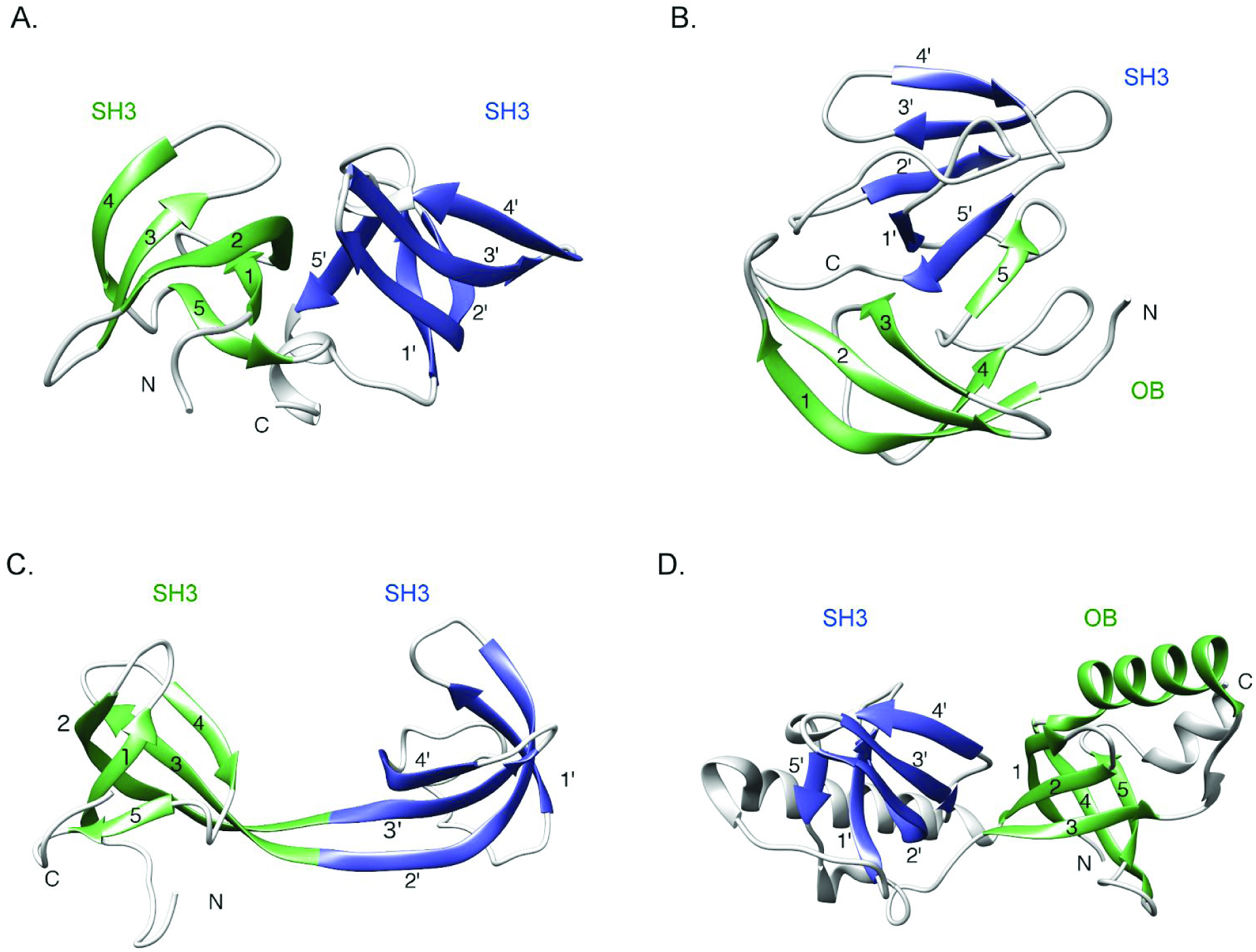
Small barrels can be combined in various ways. (A) SH3-SH3 tandem barrel in 53BP1 (PDB ID: 1SSF). (B) OB-SH3 tandem barrel in ribosomal L2 (PDB ID: 1S72). (C) SH3-SH3 interdigitated barrel in JMJD2A (PDB ID: 2QQR). (D) SH3 barrel embedded within OB barrel in TDRD (eTud) (PDB ID: 30MC). The barrel closer to the N-term - whether it is a complete (A,B) or partial (C, D) is colored in green, the second barrel is colored in blue. The β-strands in SH3 and OB are labeled using nomenclature from Figs 1 and 5, respectively.

##### 2.5.1.2 OB〰SH3 hybrid and OB〰OB tandem barrels

The OB and SH3 domains can combine in tandem in either order, as demonstrated by the ribosomal protein L2 and eukaryotic translational elongation factor eIF5A (Nakagawa et al. 1999; Dever, Gutierrez, and Shin 2014). In the L2 protein (Fig 7B)—thought to be one of the oldest ribosomal proteins (Harish and Caetano-Anollés 2012)—an N-terminal OB is connected to the SH3-like domain by a 3_10_ helix which parallels the 3_10_ helix between β4 and β5 in the SH3-like domain. The β5 strands from two barrels are arranged in antiparallel manner, thereby extending the OB sheet (Nakagawa et al. 1999). Conversely, in eIF5A the N-terminal SH3 module is followed by an OB domain (Dever, Gutierrez, and Shin 2014). Ribosomal protein S1 provides an example of six tandem OB domains, two of which are involved in binding the 30S ribosomal subunit (Demo et al. 2017; Giraud et al. 2015).

##### 2.5.1.3 SH3〰SH3: Interdigitated Tudors

The most intricate contacts between two adjacent barrels occurs in the interdigitated Tudors, such as JMJD2A (Fig 7C) (Y. Huang et al. 2006) and RBBP1 (Gong et al. 2014). These structures have been described as two barrels ‘swapping’ some strands, resulting in what has been termed an ‘interdigitated barrel’. In these cases, the long β2 and β3 strands contribute to the sheet in their parent barrel and then traverse to the adjacent barrel, yielding two compact structures wherein the first two strands belong to one ‘linear’ (in sequence) barrel and the other two strands belong to the other ‘linear’ barrel. An antiparallel β-sheet forms along the entire length of β2-β3-β2’-β3’.

##### 2.5.1.4 An SH3 barrel embedded in an OB

Staphylococcal nuclease domain-containing protein 1 (SND1) contains five tandem OB-fold domains with an SH3-like (Tudor) domain inserted into the L23 (Distal loop-equivalent) of the fifth OB barrel (Liu et al. 2010). Such an arrangement of OB and Tudor units is typically referred to as an extended Tudor domain (eTudor or eTud). The extended Tudor domain consists of two β-strands from the OB, the linker (containing an α-helix) and five β-strands of the SH3-like (Tudor) domain. Both parts of the split OB domain are essential for binding symmetrically dimethylated arginine (sDMA) residues often found in the C-terminal tails of these proteins. In this system, the OB-fold (SN domain) and SH-fold (Tudor domain) function as a single unit (Liu et al. 2010; Friberg et al. 2009). The *Drosophila* SND1 protein features 11 tandem extended Tudors, also known as maternal Tudors (Ren et al. 2014).

### 2.5.2 Possible interfaces in oligomeric assemblies

The β-strands of an SBB, particularly those that flank the domain, are typically of roughly equal length. This simple geometric property enables the flanking ‘edge strands’, such as β4 and β5 of the Sm fold, to laterally associate via backbone hydrogen bonds and other enthalpically favorable interactions between adjacent barrels; this capability, in turn, facilitates the assembly of SBB subunits into dimers, cyclic oligomers (Section 2.5.3), or higher-order states (Section 2.5.4). For instance, the five-stranded SBBs in most known Sm and Sm-like (e.g., Hfq) proteins self-assemble into (mostly) hexameric and heptameric rings via β4…β5’ interactions. Such interactions extend the surfaces of adjacent subunits to give broad patches that effectively define the ligand-binding properties of the faces of the toroidal disc; in Hfq and other Sm rings, these two faces, termed *proximal* and *distal*, function in RNA-binding (Mura et al. 2013). Though SBB-based ring-shaped architectures are reminiscent of β-propeller proteins (e.g., WD40 repeats), note that the geometry of the strand associations between SBB subunits is quite distinct from that of the blades in solenoidal β-propellers. Table 3 reports the combinations of β-strands that have been found to mediate assembly into dimers or oligomers, always in an antiparallel configuration.

**Table 3.**
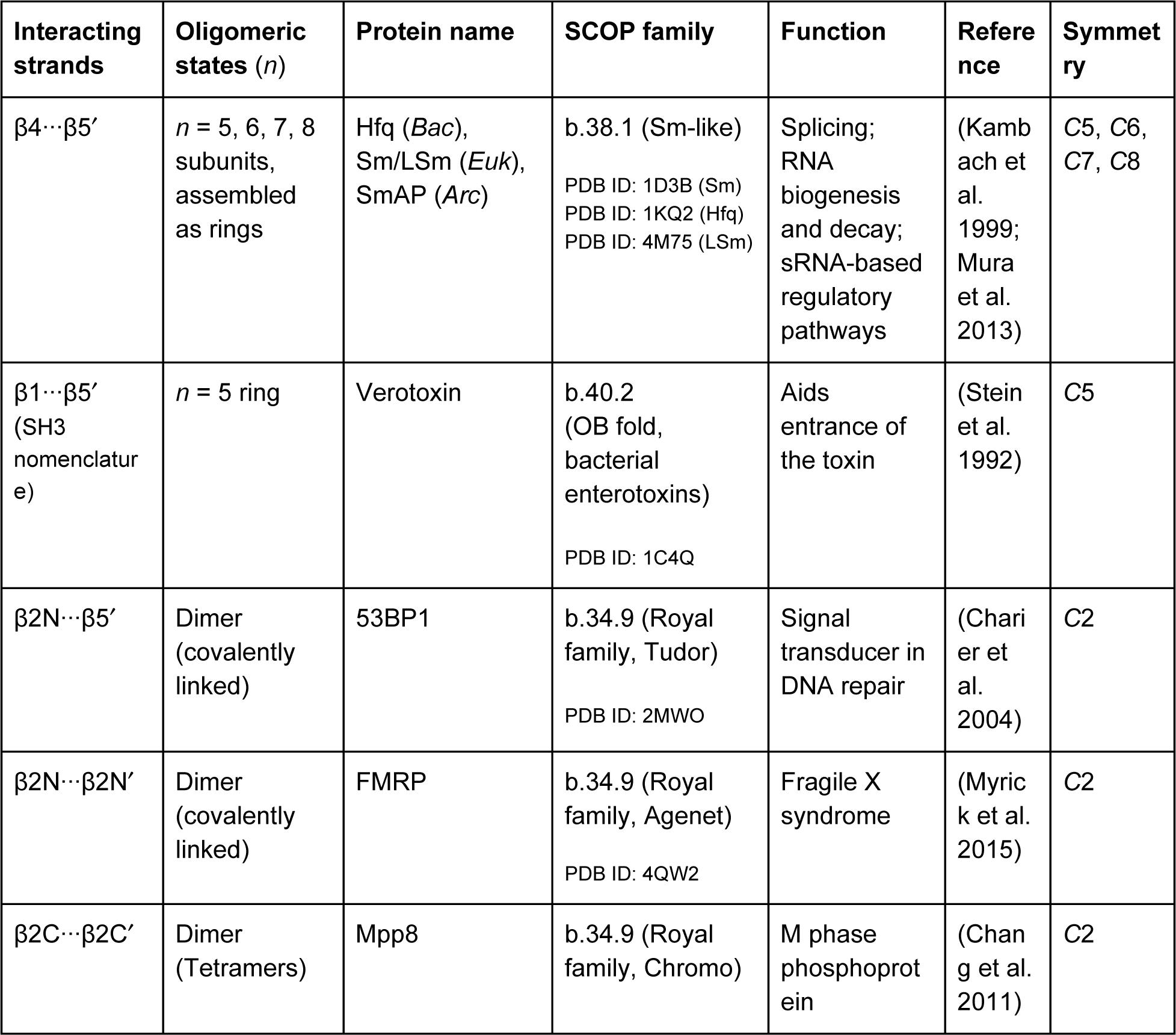
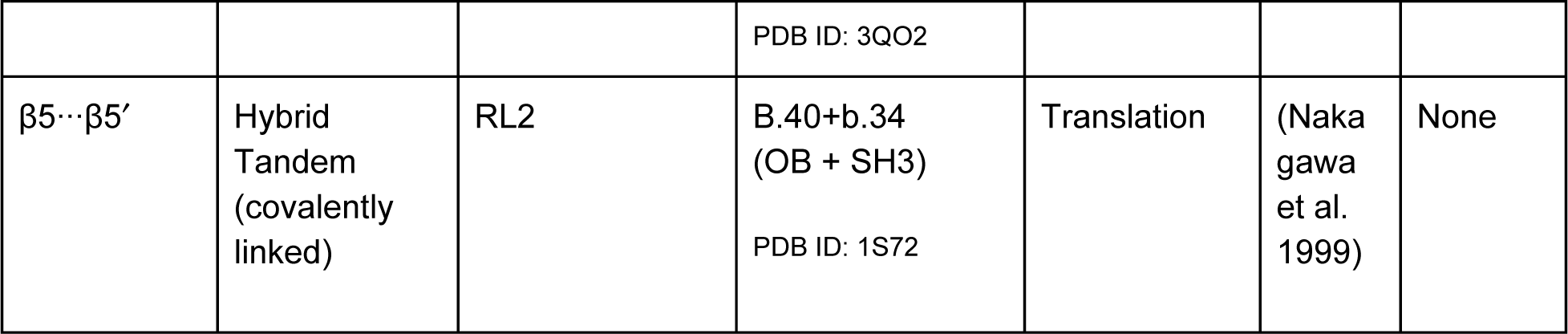
Strand-strand interactions in quaternary and pseudo-quaternary arrangements of barrels. For β2C…β5’, note that the SH3 strand nomenclature is used, even though this is an OB-fold and hence the strand numbering differs (see Table 2A for mapping). In the case of RL2, β5-β5’ hydrogen bonding in an antiparallel orientation is possible because of conformational changes in the OB domain of RL2.

Not all SBBs can oligomerize via strand…strand hydrogen bonding of the backbone. Elongation of loops, or the presence of N-or C-term decorations, often prevent the β…β interactions that mediate oligomerization. For example, the RT loop in the polyPro-binding SH3 domain (b.34.2) sterically occludes strand β4, thereby precluding β4…β5’ hydrogen bonding and ring assembly. Similarly, elongation of the n-Src loop in the case of the PAZ domain, and the N’- and C’-terminal extensions in the case of the Plus3 domain of Rft, hinder β4…B5’ hydrogen bonding. Indeed, cyclic oligomers akin to Sm rings are completely absent from the superfamily of polyPro-binding SH3 domains, though this superfamily does contain instances of dimers that form via alternative (non–β4…β5’) interfaces (Levinson, Visperas, and Kuriyan 2009); (Harkiolaki et al. 2003). Relieving obstruction of the β4/β5 strands by shortening any loops or extensions in an SBB, via mutagenesis, would be one experimental approach to test the above ideas.

Oligomerization is also possible via (i) side-chain interactions among loop residues, as in the cases of tetramer formation of HIN domains (b.40.16) (Q. Yin et al. 2013); (ii) side-chain interactions between strands, as in the dimerization of viral integrase (b.34.7) (Lutzke and Plasterk 1998; Z. Yin et al. 2016), or (iii) via α-helices present at the termini, as seen in the trimerization core of RPA (Bochkareva et al. 2002). In terms of oligomeric variability, Sm domains can assemble into pentamers, hexamers, heptamers and octamers via the classic β4…β5’ interface. In addition, they can self-assemble via alternative modes—see, for instance, an LSm4 trimer mediated by β4-β4’ interactions, discussed in (Mura et al. 2013).

By oligomerizing, an SBB domain substantially increases the solvent-accessible surface area available for the molecular interactions that typically stitch together a stable, biologically functional complex. Oligomerization also enables an SBB-based assembly to stably interact with what otherwise may have been only weakly-binding ligands (i.e., avidity). That self-assembly confers these sorts of functional advantages on a small domain, such as the SBB, is a well-recognized evolutionary process (Goodsell and Olson 2000; Ahnert et al. 2015). Extending this paradigm a step further, the assembly of different SBB homologs within a single species (i.e., SBB paralogs) can yield hetero-oligomeric complexes with novel biochemical properties, as part of an evolutionary mechanism of neofunctionalization (Veretnik et al. 2009; Scofield and Lynch 2008).

### 2.5.3 Higher-order assembly of SBBs into multimeric rings

Many single-domain SBBs self-assemble into quaternary structures that are biologically active. A well-studied example of oligomerization is the toroidal rings formed by Sm and Sm-like (LSm) proteins. The uniquely positioned β4-(3_10_)-β5 strands, which straddle the body of the barrel, lead to interactions between the β4 strand of one monomer and the β5 strand of the adjacent monomer, ultimately connecting between five and eight monomers into a doughnut-shaped ring (Fig 8A). The assembly can also be viewed as linking a three-stranded Sheet A of one monomer with a three-stranded Sheet B (Fig 1), giving a six-stranded sheet that connects the two faces of the toroidal disc.

**Figure 8.**
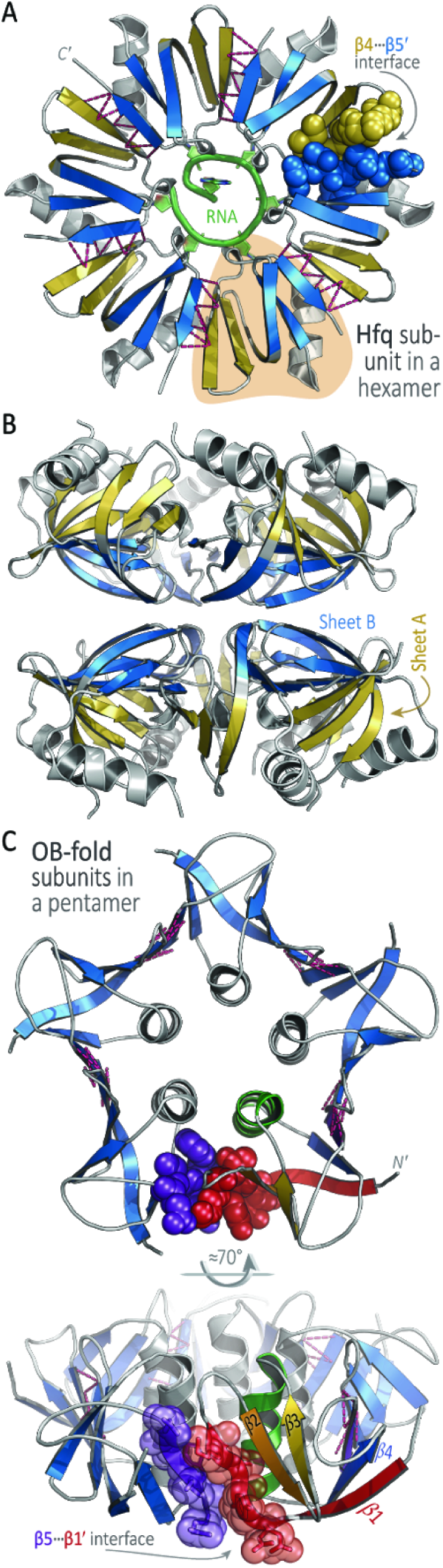
The SBB is a versatile module for oligomerization and higher-order assembly. (A) In this ribbon cartoon diagram of an Hfq hexamer (PDB 1KQ2), colored by the sheet A/B scheme (Fig 1B), one subunit is highlighted in orange (near the 5 o’clock position) and a bound RNA is drawn in green; the C’-terminus is labeled for a subunit near 11 o’clock. The crucial β4…β5’ interface generates a toroidal disc by stitching together Hfq subunits in a head-to-tail manner; these two β-strands are rendered as space-filling spheres for one subunit, and dashed magenta lines denote hydrogen bonds and other interactions between other subunits. Together, the strands of the individual SBBs effectively create a contiguous, cyclic β-sheet that comprises the body of the ring; this cyclic sheet has 30 strands in the case of Hfq (6x5), and 35 in the case of the heptameric Sm/LSm proteins (7x5). The two faces of Hfq and the other Sm rings are often found to mediate higher-order assemblies, such as in the ((Hfq)_6_)_2_ assembly shown in (B). In this panel, the same coloring scheme is used as in (A), revealing that the *distal-distal* interface of the dodecamer is built upon Sheet B of the SBB. To illustrate strand-mediated assembly in another SBB, panel (C) shows two views of a verotoxin pentamer (PDB 1C4Q), which adopts the OB fold. The β5-β1’ interface is shown as spheres and the strands of one subunit are labeled (lower-right). The β1 strand of the OB fold is structurally analogous to strand β4 of the Sm fold (see text); this example underscores the plasticity and versatility of SBBs as a structural scaffold for β-strand-mediated oligomerization.

The β4-β5 substructure of the SH3 and Sm-like fold is unique in its shape as well as its positioning with respect to the rest of the structure. Notably, this region was identified among 40 peptides that likely originated in ancient proteins (Alva, Söding, and Lupas 2015). Interestingly, we have found that structural alignment using just β4-β5 often suffices to also align the rest of an SBB.^1^ The two faces of the toroidal disc are formed by the two β sheets of individual barrels: Sheet A (*Meander*) forms the *distal* face, while Sheet B (*N-C*) forms the *proximal* face. The lateral periphery of the ring (Sauer 2013), also termed the ‘outer rim’ (Weichenrieder 2014), contains solvent-exposed residues that form the region between the *distal* and *proximal* faces. In Hfq (and not necessarily all Sm rings), the lateral rim appears to act as a site for auxiliary RNA interactions. Transiently stable RNA^1^•Hfq•RNA^2^ complexes promote annealing of the two RNA strands (RNA^1^/RNA^2^ are often an sRNA/mRNA pair), yielding a host of downstream physiological effects. The SBB residues that form the lateral site are functionally important: sRNA-binding is anchored on the *proximal* face, while the mRNA target binds mainly at the *distal* face (Sauer 2013; Weichenrieder 2014). Productive RNA interactions require the two RNA strands to physically associate, and the lateral rim of Hfq’s ring of SBBs domains appears to facilitate that process.

Several more cases of ring formation from small β-barrel proteins are known. Another example of oligomerization involving an Sm-like fold (b.38) is the bacterial mechanosensory channel MscS, formed by seven multi-domain proteins that assemble to form a pore; the central domain is a small β-barrel, which forms a heptameric ring closely resembling those of the archaeal Sm homologs (Mura, Phillips, et al. 2003; Steinbacher et al. 2007). In the case of the OB-fold (b.40), a pentameric verotoxin ring forms via β5…β1’ hydrogen bonding between monomeric subunits (Fig 8C; (Stein et al. 1992)). A final example is provided by the cell-puncturing structure in bacteriophage T4, which contains a trimer of the gp5 protein; the N-terminal domain folds as a small β-barrel (OB-fold, b.40), and forms a circular channel (Kanamaru et al. 2002).

### 2.5.4 Polymerization into fibrils and other higher-order oligomeric states

Proteins that are β-rich are prone to polymerization and formation. The resulting polymeric species, which resemble amyloidogenic ultrastructures (β-rich fibrils), may be physiologically functional in some cases, or toxic and pathogenic in other instances. The structural unit from which a fibril forms can be an individual SBB, a toroidal ring, or a double-ring assembly (with either head-head or head-tail stacking).

A common pathway to fibril formation for SH3 polyPro-binding domains begins with domain swapping between two protomers, in which any loop (RT, n-Src or Distal) can function as a hinge to partially open the β-barrel and exchange β-strands with the other protomer. In the open (swapped) state, such interacting ‘hinge loop’ regions may become rigidified and may contain short β-strands. These β-strands can then serve to nucleate further self-assembly into amyloid (Cámara-Artigas 2016). Alternatively, the hydrophobic strand (β1 or β4, for OB or SH3) may not undergo typical pairing with β5 if the latter is disordered, thereby freeing it to form non-native contacts with β1 of other protomers; such a process yields aggregation-prone intermediates that can lead to fibrillization (Neudecker et al. 2012). Both of these pathways are strongly tied to the folding process for the SBB domain itself. Mutations that destabilize folding are found primarily in the open loops/hinges and unpaired strands. This property enables non-native folds with swapped domains, as well as polymerization via β-strands and, ultimately, fibril formation.

Polymerization and fibril formation via the stacking of SBB rings has been seen with the bacterial Hfq and the Sm-like archaeal proteins (SmAP). Structurally, one route seems to proceed through stacking of either the *distal* or *proximal* faces of two rings. SmAP rings have been shown to stack *proximal-to-distal* (Mura, Kozhukhovsky, et al. 2003), while Hfq homologs have been found in *proximal-to-distal* (Stanek et al. 2017) and *distal-to-distal* (Schumacher et al. 2002) orientations. Certain geometric arrangements of Hfq and SmAP rings allow for runaway assembly into fibrillar polymers. In Archaea, these fibrils are formed by the self-assembly of multiple *proximal-to-proximal* stacked SmAP rings, yielding striated bundles of polar tubes (Arluison et al. 2006; Mura, Kozhukhovsky, et al. 2003). *E. coli* Hfq rings can also self-assemble laterally into slab-like layers, each layer built of six hexameric rings to give a 6x6 arrangement in each layer. Fibrils are then built via the stacking of such layers (Arluison et al. 2006). The C-terminal region of the 102-residue *E. coli* Hfq (comprising 30% of the protein) is intrinsically disordered and was shown to be critical for the assembly of fibrils into higher-order cellular structures (Fortas et al. 2015).

While higher-order oligomers and polymers of SBB proteins have been detected in multiple systems using different experimental approaches (crystallography, electron microscopy and image analysis, ultracentrifugation), the potential roles of such species remains enigmatic. A potential way in which SBB-based fibrils may be of (non-pathogenic) relevance, *in vivo*, is as a concentration-dependent molecular switch. Namely, at low protein concentrations an Hfq or Sm ring would form and go about its ‘normal’ (constitutively active) cellular function in, say, some RNA-associated pathway; this corresponds to the ‘on’ position of the switch. At high concentrations, polymerization of an Hfq or Sm protein ring into fibrils would sequester its RNA-binding sites (e.g., the *proximal* face), thus silencing the protein’s RNA-related activity.

### 2.6. Folding and stability

#### 2.6.1 Folding of small β-barrels

Folding of the SBB domain seems exceptionally robust—the same fold is achieved by a wide range of sequences, and also when the sequence elements are permuted (Fig 2) or mutated. Detailed folding studies have been performed with polyproline-binding SH3 domains (b.34.2) and on the OB domains in cold-shock proteins (b.40.4.5). For SH3 domains, folding proceeds via two-state kinetics, i.e., an infinitely cooperative *unfolded* (U) ⇌ *folded* (F) transition. The high-energy transition state is characterized by multiple conformations of partially-collapsed structures, and is termed the transition state ensemble (TSE). It has been consistently found that the partially folded conformational states (i.e., the TSE) of SH3 and OB folds are bipartite: they contain (i) a hydrophobic region that nucleates further folding, consisting of most of the β2-β3-β4 segment (i.e., *Sheet A* or the *Meander*, β2-ββ-β4), and (ii) conversely, a *Sheet B* (or *N-C*), which includes β1+β5, and which is disordered in the TSE (Chu et al. 2013; Neudecker et al. 2012; Riddle et al. 1999; Viguera, Blanco, and Serrano 1995). For the OB fold in cold-shock proteins (CSPA, CSPB), an intermediate state was recently proposed; it, too, consists of the three-stranded β-sheet, β1-β2-β3, which structurally corresponds (Fig 5) to the *Meander* of the SH3 fold (L. Huang and Shakhnovich 2012).

The robustness of the folding process is largely attributable to its cooperativity, which stresses the significance of local interactions during folding: residues that initiate the folding process are local in sequence (Riddle et al. 1999; Martínez and Serrano 1999; Baker 2000). The hydrophobic zipper (HZ) model of Dill and colleagues (Dill, Fiebig, and Chan 1993), which begins with local interactions and eventually brings together more distant residues to form the hydrophobic core via β-hairpin formation, is consistent with what is known about small β-barrels. The HZ substructure is formed by a group of neighboring residues (cooperativity), thereby relaxing the need for specific residues (tertiary contacts) to achieve folding. Indeed, formation of a three-stranded meander in SH3 domains and in the small, modular WW domain—both of which are well-studied protein folding systems (Riddle et al. 1999)—always begins with one or both of the loops/turns (Davis and Dyer 2016; Jager et al. 2008; Maisuradze et al. 2015). The folding of CspA/CspB also initiates within the loops, as would be expected were interactions among the local residues responsible for driving the folding process (L. Huang and Shakhnovich 2012; Vu, Brewer, and Dyer 2012).

The significance of local interactions is also supported by circular permutation engineering within the SH3 domain of alpha-spectrin (Viguera, Blanco, and Serrano 1995); (Martínez et al. 1999). In such experiments, the N-and C-termini are covalently stitched together and the polypeptide sequence is cut open by introducing N/C termini in one of the three loops, thus rearranging the sequential order of secondary structures. Intriguingly, all such permuted constructs were found to adopt the same fold; however, the order of folding differed, as the β-hairpin formed by the linked ends (β1–β5) appeared early in the folding process.

Several SBB-containing structures have entire domains embedded into one of the loops of the barrel. In the cases of extended Tudors, the 96-amino acid, SH3-like Tudor domain is inserted between β2 and β3 of an OB fold (Friberg et al. 2009). In the case of BRCA2, the 154-residue Tower domain occurs between β1 and β2 of the OB domain (Yang et al. 2002). Finally, there also exists a signal peptidase (SPase; a rare case of enzymatic activity within an SBB) wherein an ≈110-amino acid region, comprising residues 150-266, is inserted between strands β2 and β3 (n-Src loop) of the SH3-like fold (Paetzel et al, 2002); notably, that particular structure (PDB 1B12) tests the limit of what one would classify as a *bona fide* SBB, as the β4 strand is very short, there are scant β2…β3 interactions, a helical turn links β3-β4, and only a short linker (not a short 3_10_ helix) lies between β4〰β5. All known cases of loop insertions occur in the *Meander* sheet, meaning that this sheet can tolerate significant distances (in sequence) between its constituent strands; this, in turn, implies that non-local interactions also can suffice to nucleate *Meander* sheet formation, in lieu of what otherwise occur as local interactions in SBBs without insertions (e.g., between adjacent strands in a β-hairpin). An intriguing open question concerns the relative contributions of local and non-local interactions in shaping the free energy landscape for SBB folding, including nucleation of nascent (sub)structures such as the *Meander* sheet. This question could be experimentally probed via protein engineering, biophysical characterization, and structural approaches. For instance, would a mutant SPase, wherein the >100-residue β2〰β3 insertion were excised and the (n-Src) loop closed, fold with similar thermodynamic and kinetic properties as the wild-type protein? What about if the lengths of the loop insertions were systematically varied? Also, would such engineered constructs adopt the same 3D structures (particularly in the *Meander* region)? Pursuit of these types of questions would illuminate the underlying mechanisms by which SBBs fold into stable structures, including the relative roles of local and non-local interactions.

Perhaps the most poignant evidence of the resilience of the SBB structure, as concerns folding, is the unusual case of RfaH. In RfaH, the C-term domain spontaneously switches from an α-hairpin structure (when bound to the N-term domain) to the small β-barrel structure (when released from interaction with its N-term domain). Such a change in structure has far-reaching functional consequences; remarkably, RfaH plays key roles both in transcriptional elongation and in translation initiation (Burmann et al. 2012).

#### 2.6.2 Structural stability, resistance to thermal and chemical denaturation

The compactness and robustness of the SBB fold has ramifications for the stability of SBB-containing proteins. Experimental studies of Sm, LSm and Hfq homologs have demonstrated that these SBB proteins resist unfolding by thermal or chemical denaturation. For instance, samples of Hfq homologs (even from mesophilic species, such as *E. coli*) can typically be heated to 70-80 °C for 10-20 minutes without denaturation or loss of solubility (Zhang et al. 2002; Stanek and Mura 2018). Similar resistance to thermal denaturation has been found in the SH3-fold family of tyrosine kinases (Knapp et al. 1998), and has long been known to occur with SH3 domains more generally (e.g., an SH3 domain from the soil-dwelling nematode C. *elegans* was found to melt at T_m_ ≅ 80 °C (Lim, Fox, and Richards 1994)). The OB-fold-containing verotoxin from *E. coli* exhibits moderate thermostability and retains activity even after 10 minutes of heating to 60 °C (Yutsudo et al. 1987), while an OB-fold protein from the mesophilic (and radiation-resistant) bacterium *Deinococcus radiopugnans* has a 30-minute half-life at 100 °C (Filipkowski, Koziatek, and Kur 2006). The structural and physicochemical basis for the thermostability of SBBs has not been elucidated, although, as mentioned above, studies on the SH3 domain suggest that local interactions involving the HZ substructure are likely important determinants of stability. More broadly, the SBB fold’s resistance to thermal and chemical denaturation makes it a promising module in protein engineering and design efforts.

## 3. Conclusions

The small β-barrel (SBB) domain pervades much of biology, including nucleic acid-related pathways (e.g., RNA metabolism, DNA maintenance, ribosome assembly and RNA-based regulatory circuits) as well as other, entirely disparate, milieus (e.g., membrane channels). SBB-containing protein families occur across the tree of life, with many representatives conserved in archaeal, bacterial and eukaryotic lineages. The ancient SBB fold likely arose in ribosomal proteins and it appears to have been recruited extensively, over the aeons, to serve myriad functional roles. The SBB domain often acts as a structural platform that scaffolds the assembly of eukaryotic ribonucleoprotein complexes, as a protein module in signal transduction pathways, and as a chaperone of RNA…RNA interactions in bacterial sRNA-mediated regulatory circuits. The lack of a distinct, reliable sequence signature has hampered the identification of SBB proteins via sequence similarity searches; thus, the fractions of SBBs in various genomes, as well as the full breadth of their functional repertoire, remain unknown. The range of SBB functions described in this work—broad though it may seem—still represents only a subset of known functionalities. Many SCOP folds can be classified as members of an SBB superfold, and we limited the scope of this work to those select superfamilies that are highly represented in the structural and bioinformatic databases (SH3, Sm, OB).

The SBB ‘fold’ is really a ‘superfold’, insofar as it encompasses several fold families (SH3, Sm, OB, etc.) that may be non-homologous. Because even just one of these fold families includes a vast swath of biochemistry and cell biology, historically much emphasis has been on the unique properties of a particular subset of SBB proteins (e.g., Sm proteins and their roles in snRNP cores), rather than on uncovering any unifying principles. That is, any parallels between the many SBB-containing protein families have been lost in a sea of idiosyncrasies for each of the various families, so any recurring themes have gone largely unrecognized. To help identify recurrent themes and patterns, this work has sought to systematically define and survey the SBB domain, chiefly in terms of structure ↔ function relationships, and their evolutionary contexts. An initial step has involved terminology: as in many areas of science, alternative (and sometimes incongruous) descriptive schemes and nomenclature systems have emerged for describing closely related entities and the basic relationships between those entities (e.g., for Sm and SH3 folds). Thus, in this treatment of SBBs we have identified alternative nomenclatures and mapped them to one another as much as possible. More broadly, protein domains such as the SBB challenge us to develop systematic, formalized structural description frameworks that can transcend the SCOP, CATH (Dawson et al. 2017), and ECOD classification systems (Cheng et al. 2014) in order to accommodate (and precisely ‘capture’) the deep structural and functional plasticity of the SBB and SBB-like superfolds.

A hallmark of the SBB is its marked variability, in terms of the 3D structures of individual domains, its known oligomeric states and higher-order quaternary structures, and its overall functional plasticity (types of cellular pathways; RNA-, DNA-and protein-binding capacities). Is there any specific set of principles—any salient structural, physicochemical, dynamical properties—that account for such deep variation? How does the SBB achieve such functional versatility, while maintaining a stable, unique structural framework that defines it as a discrete superfold (distinct from its neighbors in fold space)? In elucidating SBB sequence ↔ structure ↔ function relationships, a key issue is that residue positions which compose different regions of an SBB surface (e.g., ligand-binding patch on an Hfq ring) contribute quite differently to functional properties (e.g., RNA-binding specificity), in comparing one type of SBB to another (e.g., Hfq versus Tudor); notably, such is the case despite the great structural similarity between the domains. Indeed, the fact that the SBB structure is preserved even when the strand order is permuted (i.e., SH3 versus OB superfamilies), implies that this domain architecture is a resilient platform for deep sequence variation (and, thus, variation in function). In this way, the SBB challenges our usual perspective of sequence variation yielding concomitant variation in structure↔function relationships. The SBB fold’s ability to accommodate profound structural variation also raises intriguing questions about whether there exist well-defined boundaries of the SBB in fold space and, assuming so, what its nearest structural neighbors might be. Are there other small β-barrels that are structurally distinct from the SBBs defined here? These are open questions.

**One emerging theme** from our analysis is that the SBB’s β-barrel is a robustly-folding, compact structure for elaborating new biochemical functionality (a wide range of sequence space can apparently adopt the SBB fold). For example, an SBB’s electrostatic properties can be altered by its set of solvent-exposed residues, thus affording a means to finely tune interactions with nucleic acids. The termini of the SBB often vary greatly, in terms of the presence/absence of helices or other secondary structural elements, and the loops of the SBB also vary immensely—both in sequence and in length (from a tight β-turn to the insertion of entire functional modules/domains). Apparently, the extent of possible sequence and structure variation, and hence the range of potential interactions with RNA, DNA, proteins and ligands, is immense.

**A second emerging theme** is the tendency of SBB-containing proteins to oligomerize into biologically functional units. The molecular contacts that stitch together such assemblies are often mediated by the edge strands flanking the SBB fold. Oligomerization confers many benefits, in terms of biochemical functionality. For instance, self-assembly into homomeric complexes affords much greater surface area for binding to other biomolecules (e.g., in Hfq-RNA interactions), while the assembly of SBB domains into heteromeric complexes yields the further advantage of enabling asymmetric assemblies to form (e.g., the seven Sm paralogs that nucleate the hetero-heptameric snRNP core complex). Indeed, the oligomeric plasticity and quaternary structural diversity of the SBB domain may well distinguish it among all known protein folds.

**Finally, a third emerging theme** is the severe modularity of the SBB in most SBB-containing protein families. The ≈60-residue SBB domain occurs throughout the proteome, often as part of a larger polypeptide that extends the SBB core in the N-or C-terminal directions (by just a few residues, or even entire domains). Similarly, anywhere from a few amino acids to >100 residues have been found inserted into the loops of an otherwise intact, canonical SBB domain, and some proteins contain tandem repeats of SBB domains (echoing the behavior of RRM-containing proteins). Such extensive modularity is evolutionarily adaptive. The SBB may be the only known fold that functions robustly, and broadly, in three contexts: on its own (as a monomer), as a structural unit in quaternary assemblies, and as a domain within multi-domain proteins.

A detailed functional analysis of SBBs could have, as one aim, elucidation of the structural mechanisms by which SBBs recognize different classes of targets (e.g., OB…ssDNA versus OB…protein binding). Notably, the RRM domain at least superficially resembles SSBs, in terms of structure/function relationships: the RRM is a small, four-stranded antiparallel β-sheet (with helices at both termini), it binds RNA (as do many SSB proteins), it exhibits a great degree of structural variation, and it is functionally quite versatile (interacting with a wide variety of possible ligands, including RNA, DNA and other proteins). Determining the fundamental structural and physicochemical principles that enable the deep structural and functional plasticity of SSBs, RRMs, and other β-rich fold families represents a broadly stimulating area for future work.

## Acknowledgements

Portions of this work were supported by the Jeffress Memorial Trust (J-971; CM) and NSF CAREER award MCB-1350957 (CM).

## Footnotes

1 This implies that whatever structural role is served by the β4/β5 pair (e.g., oligomerization, in Sm/Hfq proteins) may be a significant constraint on the evolutionary drift of those residue positions that dictate the relative geometric disposition of strands β4 and β5.

